# Growth rate dependent DNA methylation patterns along bacterial chromosomes

**DOI:** 10.64898/2026.05.23.720832

**Authors:** Akshat Mall, Mohammad H. Abbaspour, Dawson J. Mathes, Klas I. Udekwu, Christopher J. Marx

## Abstract

DNA methylation plays critical roles in gene regulation in bacteria, from regulating essential processes like the cell cycle to phenotypes of practical interest like pathogenicity and motility. Synthetic manipulation of global methylation levels has broad impacts on cellular physiology, changing expression patterns of hundreds of genes. However, whether or how environmental variation in natural settings similarly impacts DNA methylation patterns has been unclear. In this work, using the alphaproteobacteria *Methylobacterium extorquens* and *Caulobacter crescentus* as model systems, we discover the methylome is highly fluid in response to environmental variation, with different environments leading to distinct patterns of increased or decreased methylation levels along the chromosome. Despite a heterogeneous effect of different environments on methylation patterns, we find a general principle where the dependence of methylation states on position in the genome decreases in proportion to growth rate. A simple model that considers the methylation state through different phases of the cell cycle as a function of distance from an origin provides a framework to interpret the effects of different stressors upon the observed environmental responsiveness of the methylation patterns. Our work highlights how sequencing data alone can shed light on important aspects of microbial physiology.

**Significance Statement:** DNA methylation is known to profoundly impact gene regulation in prokaryotes, with both distinct methylation states at specific loci and global levels of DNA methylation modulating critical cellular phenotypes. Yet whether or how DNA methylation patterns depend on environmental variation remains unclear. Using *Methylobacterium* and *Caulobacter* as model systems, we combine experiments and theory to uncover general principles of how global patterns of DNA methylation are shaped by the environment. In particular, we discover a positive relationship between growth rate and the magnitude of the genomic position-dependent methylation level, highlighting how sequencing data can provide a culture-independent approach to estimating microbial traits like growth rate in natural settings. Our results resolve an open question, highlighting that DNA methylation patterns in bacteria can rapidly change in response to environmental shifts and revealing rules by which methylation patterns can help understand cellular phenotypes.

## Introduction

DNA methylation, in recent years, has garnered significant attention in prokaryotic systems^1^, where it serves functions as part of restriction-modification systems^2^, DNA mismatch repair^3^, and regulating gene expression^4,5^. In bacteria, two adenine methyltransferases – Dam in gammaproteobacteria (targeting GATC sites), and CcrM in alphaproteobacteria (targeting GANTC sites) – have been the subject of most work in understanding the role of DNA methylation in regulation of gene expression. For both clades, several genes are known where the methylation state of one or more sites in the promoter changes transcription factor affinity, and consequently expression levels^6–9^, notably virulence and motility associated genes in pathogens. Additionally, several global transcriptional regulators have been identified which show different affinities for different methylation states (fully-methylated, hemi-methylated, or unmethylated) at the target site^10–12^. Genetic mutants changing the expression of methyltransferase genes have further discovered hundreds of genes whose expression levels depend directly or indirectly on global levels of DNA methylation^13–15^.

Such modulation of cellular phenotype via changes in DNA methylation is known to serve two functions in bacteria. First, in cell-cycle regulation, where DNA replication changes methylation states from fully-methylated in the old strands to hemi-methylated in the newly synthesized strands, resulting in changing methylation and consequently, gene expression patterns which guide progression of the cell cycle^16^. This role is particularly prominent in alphaproteobacteria where CcrM expression is cell cycle regulated, transcribed only transiently when the *ccrM* promoter is hemi-methylated after passage of the replication fork^17^. These dynamics lead to a chromosome position dependence of DNA methylation patterns where sites close to the origin of replication are hemi-methylated for long periods, until CcrM is active, while sites close to the terminus remain fully-methylated^18^. Modifying the distance from the origin for a gene (under epigenetic regulation) has been shown to change the average methylation state and alter gene expression patterns^11^. Second, stochastic heterogeneity in methylation patterns leads to phenotypic heterogeneity in isogenic populations which helps maximize population fitness in complex (via division of labor)^19^ or unpredictably fluctuating environments (via bet-hedging)^20–22^.

While the prominent role of DNA methylation on gene expression has been established, it remains unclear how fluid DNA methylation patterns are – whether or how they respond to environmental variations. Recently, some environments were shown to change average methylation levels across the *Escherichia coli* genome^23^, and phosphate limitation was shown to change local methylation levels and gene expression for some *Caulobacter crescentus* loci^24^. These results suggest methylation patterns can also change rapidly in response to environmental cues but general principles relating methylation changes to environments are lacking.

Additionally, recent advances in methylome analyses due to long-read sequencing technology may provide a window into microbial function that is complementary to metagenomics and other omics strategies. In this regard, it may be similar to the use of sequencing coverage bias along chromosomes that, at least for fast-growing microbes, has been suggested to reflect differences in growth rate of microbes in natural microbiomes^25,26^.

In this work, we primarily use the alphaproteobacterium *Methylobacterium extorquens* to ask how fluid global methylation patterns are with changing environments and rules governing such patterns. *Methylobacterium* strains are commonly associated with leaf and soil microbiomes^27,28^, playing roles in influencing plant growth^29^, carbon cycling in the environment^30^, and biotechnological applications^31,32^. Using nanopore sequencing, we characterize the methylome of *M. extorquens* and find the methylation levels are quite varied across methylation sites, yet in ways that are very consistent. By capturing positional and compositional variation, a vastly cleaner picture emerges that exposes a methylome that is highly flexible in response to environmental variation. Notably, these patterns in the methylome change along the chromosome in a predictable manner according to growth rate, but with unique aspects consistent with different stressors affecting different phases of the cell cycle. We confirm the generality of the growth rate dependence of global methylation along chromosomes using both *C. crescentus* and *E. coli* as alternate model systems. A simple theoretical model based on dynamics of cellular growth phases provides a framework in which to interpret the observed shifts in methylation patterns.

## Results

### Methylation motifs in *Methylobacterium*

We first attempted to identify the DNA motifs which are frequently methylated in *Methylobacterium extorquens* PA1. Using nanopore sequencing (Methods), we discovered two frequently methylated motifs, GANTC and GCCGATC, with the adenine nucleotide in both motifs harboring 6mA modifications. GANTC motifs have been shown to be involved in epigenetic gene regulation across alphaproteobacteria, and are targeted by a highly conserved orphan methyltransferase, CcrM. Using the REBASE database^33^, we identified putative DNA methyltransferase genes in the genome linked with both of these modifications (Table S1). Given the conserved essential role of CcrM in gene regulation, we only consider GANTC sites for further analysis in this work. The number of GANTC sites in the *M. extorquens* genome was significantly lower than expected by chance (∼60% lower, Supplementary Methods), particularly in coding regions. This biased distribution of GANTC sites in the genome was similar across a diverse set of alphaproteobacteria (Fig S1A, Table S2). This was in stark contrast to the well-studied GATC motifs in gammaproteobacteria which occur as expected within genes and are particularly underrepresented in intergenic regions (Fig S1B, Table S2). This suggests distinct evolutionary forces acting on methylation motifs in different bacterial clades. The distribution of these nucleotide motifs where they do not function as targets of methylation (i.e., GANTC in gammaproteobacteria and GATC in alphaproteobacteria, Fig S1) did not show these biases.

### Global methylation patterns in *Methylobacterium* identify two putative replication origins

To understand the global patterns of DNA methylation, we first analyzed the methylome of both *C. crescentus* and *M. extorquens* in exponential phase, finding most GANTC sites were highly methylated, but with wide variance in the level observed for each site (Fig 1A,C). Despite these wide distributions, there was great consistency between replicate samples (Fig 1B,D), suggesting that this does not represent measurement noise, but rather either measurement bias or true biological differences in average methylation states for different sites. To search for potential biological underpinnings of this variance, we first examined the role of genomic position on methylation patterns. A unimodal relationship between average methylation state of a site and position on the chromosome is expected for *C. crescentus*^18^, with the replication terminus corresponding to the region with highest fraction of sites being methylated, and origin of replication corresponding to the lowest. This is because of the cell cycle dependent dynamics of CcrM activity where sites closer to the replication origin wait longer before CcrM is active, compared to sites closer to the replication terminus. For *C. crescentus*, we observed the expected unimodal, V-shaped dependence of DNA methylation levels on chromosomal position with sites close to the origin being less methylated (Fig 1E). However, the chromosome position dependent pattern of DNA methylation for *M. extorquens* represented a W-shape (Fig 1F), suggesting the presence of two origins of replication in *M. extorquens* corresponding to each of the two minima, while the replication termini correspond to the maxima (Fig 1G illustrates a model of the distinct dynamics for *C. crescentus* and *M. extorquens*).

**Fig 1.**
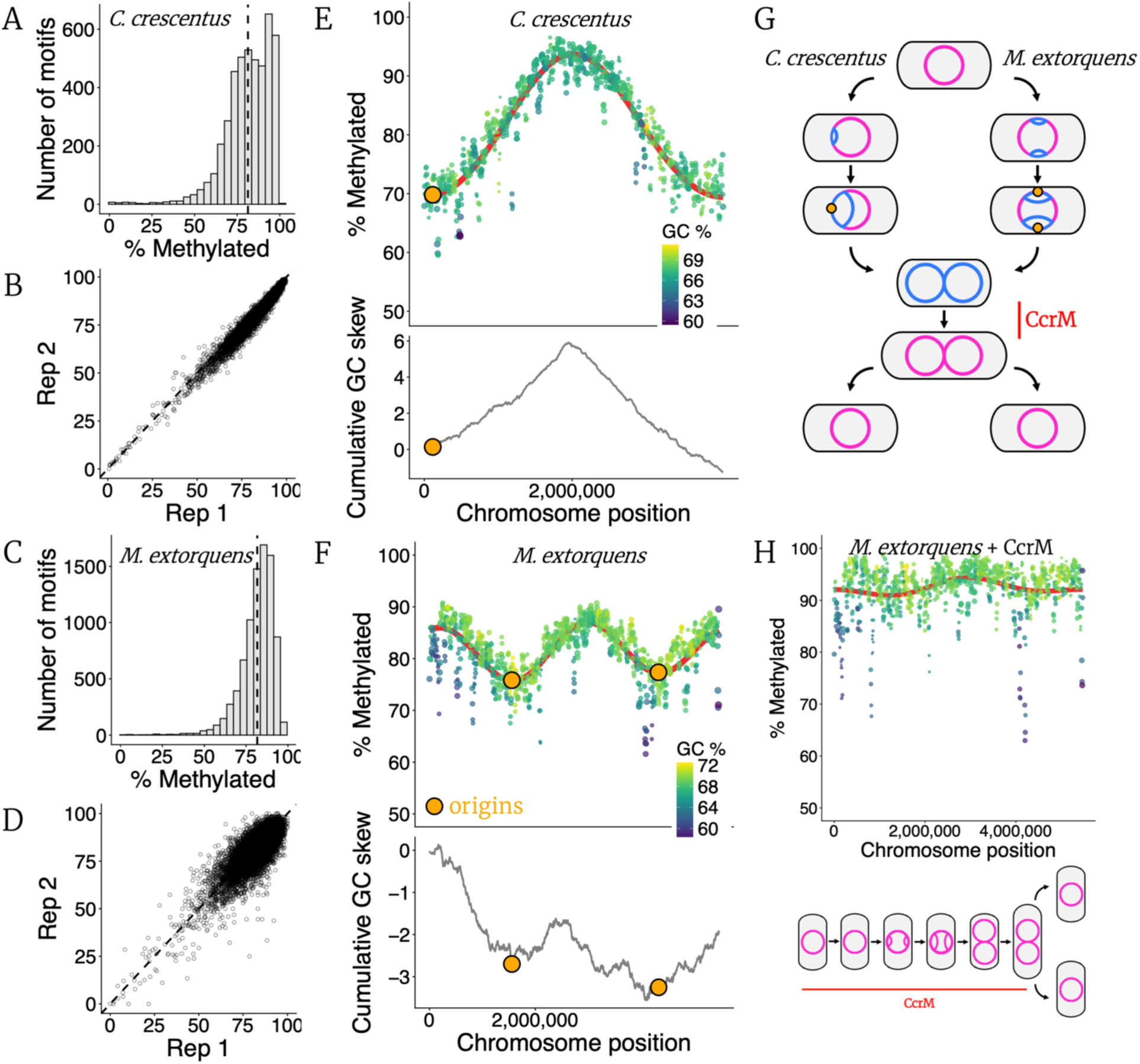
Global methylation patterns identify two origins of replication in *M. extorquens*. A &. **C)** Distribution of number of sites which are seen methylated a particular fraction of times for GANTC sites in *C. crescentus* growing on PYE (A) and *M. extorquens* growing on succinate (C). **B & D)** Scatter plot of percent methylation for each GANTC site in the genome between replicate populations of *C. crescentus* (B) and *M. extorquens* (D). **E & F)** Top panels – Chromosome position dependent pattern of DNA methylation for GANTC sites in *C. crescentus* (E) and *M. extorquens* (F). Each point represents average proportion of methylated reads for all GANTC sites in a 10,000 bp window around that point (points separated by 5000 bp). Size of points are proportional to number of GANTC sites in that window. Red curves denote best fit from a Fourier regression (Methods). Points are colored based on GC content of the 10,000 bp window around that point (darker colors denote lower GC content). Orange points represent putative origins of replication. Bottom panels – Cumulative GC skew (Methods) for *C. crescentus* and *M. extorquens* respectively, identifying the same pattern for replication origins and termini. **G)** Conceptual model of the replication dynamics giving rise to the position dependent DNA methylation patterns for GANTC sites in *C. crescentus* (left) and *M. extorquens* (right). At the beginning of the cell-cycle, all sites in the genome (circle) are fully-methylated (pink). As DNA replication begins, new strands are hemi-methylated (blue) and remain so until CcrM activity (red) at the end of the cell-cycle when the new strands are remethylated (pink). **H)** DNA methylation patterns for GANTC sites in *M. extorquens* with constitutive CcrM expression (red line throughout the cell cycle in bottom panel).

Two alternative approaches corroborate the inference of two replication origins in *M. extorquens*. First, sequencing coverage across the genome showed a similar position dependent pattern with sites near the origin overrepresented in observed reads relative to sites near the terminus (Fig S2). However, the strength of this pattern was much weaker (𝑅^2^ = 0.27) relative to the position dependent pattern for methylation (𝑅^2^ = 0.50). Second, cumulative GC skew^34,35^ independently identified the same pattern of a single origin in *C. crescentus* (Fig 1E, Methods) and two replication origins and termini in *M. extorquens* (Fig 1F). This signature of multiple origins of replication was also present in some other *Methylobacterium* strains, and in *Vibrio cholerae* strains where the function of two origins has been verified^36–38^ (Fig S3).

To directly test the hypothesis that the position dependence of methylation in *M. extorquens* was due to the late, short window of CcrM expression during the cell cycle, we cloned and expressed the *ccrM* gene using an inducible promoter. Constitutive induction of CcrM increased the total methylation and nearly eradicated its position dependence along the chromosome (Fig 1H), highlighting the dependence of this pattern on the dynamics of CcrM activity. The chromosome position dependent pattern of DNA methylation was also observed during growth on other substrates like methanol (Fig S4A), but not for stationary phase cells (Fig S4B), or for GCCGATC motifs (Fig S4C). The dependence of DNA methylation patterns on chromosome position was also greatly diminished for *E. coli* where Dam methylase is expressed constitutively (Fig S5).

### Regions with low GC content have decreased average methylation levels

While chromosome position explained much of the variance in average methylation states, we observed that replicate populations still exhibited consistent deviations from the expected methylation state based on chromosome position (Fig S6A). In particular, several ‘icicles’ of consecutive regions of abnormally low methylation were seen along the genome. We then investigated the role of nucleotide composition neighboring the GANTC sites in the genome, which revealed a strong influence of the neighboring local GC %. This signal was strongest for ∼100 bp windows around a GANTC site (Fig S6B) and explained much (𝑅^2^ = 0.44) of the remaining deviation from expected methylation percentage at that chromosomal position (Fig S6C). These influential windows are much larger than the 10-nucleotide contexts in which nanopore signals are read, making measurement bias unlikely. This suggests hypomethylation may be due to either biases in CcrM activity^39^ or inhibited access of CcrM to those regions due to competition with specific DNA binding proteins which target AT-rich sequences like certain transcription factors^40^ or more broadly with nucleoid associated proteins^41^.

### Methylome changes with changing substrate

To test how the observed patterns of DNA methylation change with changing environments, we first compared the methylation pattern of *M. extorquens* when growing exponentially on either of two substrates – succinate or methanol. Between the two carbon sources, most GANTC sites retained the same methylation state, but a few sites showed a significant change in methylation pattern, with the fraction of methylated reads decreasing or increasing significantly (Fig 2A, Table S4). The strongest changes involved flipping of the methylation state of two GANTC sites upstream of an operon encoding for one of the formate dehydrogenase gene clusters (Fig 2B) that encodes a key enzyme for methylotrophic growth^42^. As a control, no significant changes in methylation patterns were observed for replicate treatments of either succinate or methanol (Fig S7A,B).

**Fig 2.**
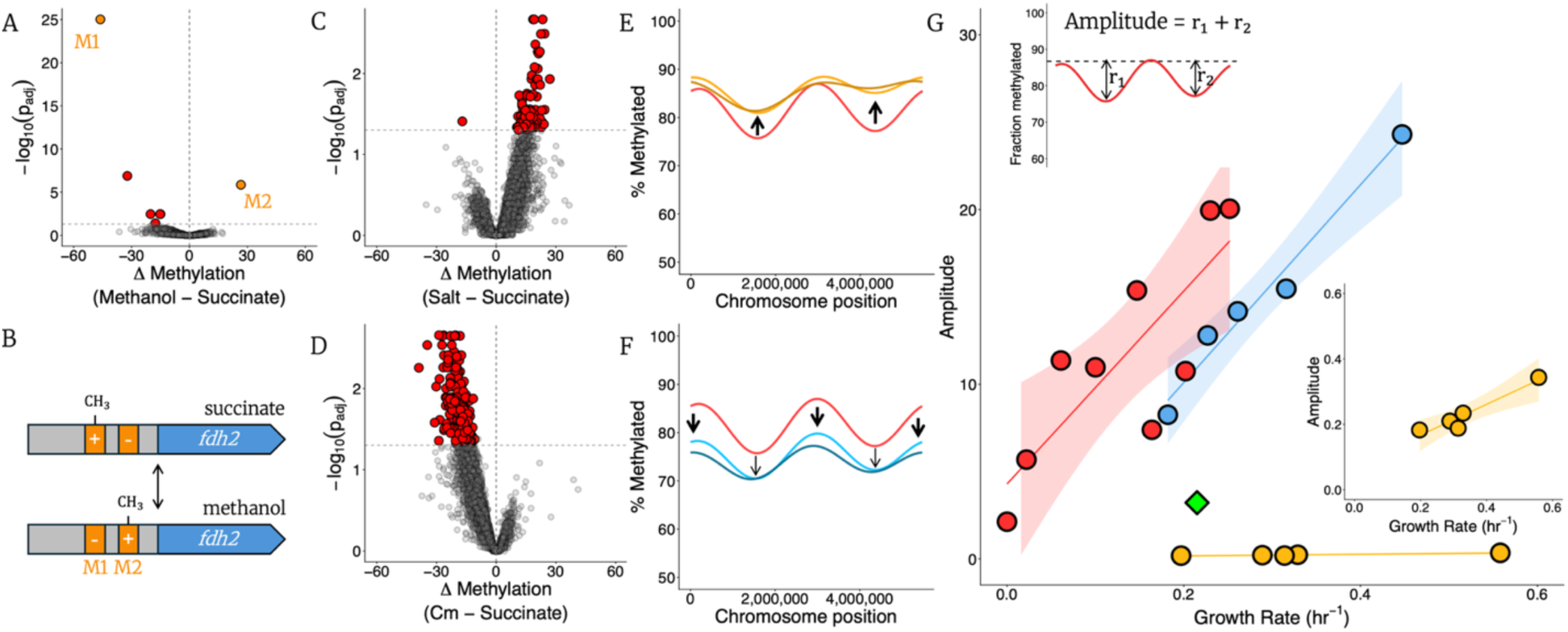
Environments change methylation patterns in a growth-rate dependent manner. **A)** Change in methylation states for populations growing on methanol compared to succinate. Each point represents the observed change in proportion of methylation reads (x-axis) for one GANTC site and the *p*-value associated with the change (y-axis). Few GANTC sites exhibit a statistically significant shift in methylation pattern (colored circles) while most sites retain the same methylation state (gray circles). **B)** The biggest methylation changes for cells growing on methanol relative to succinate are observed for two GANTC sites (orange) in the promoter region (gray) of formate dehydrogenase (blue, Mext_4404), with one site (M1) flipping from + (methylated) to – (unmethylated) and the other site (M2) doing the reverse. The two sites are located 110 and 159 bp upstream of the coding sequence. **C & D)** Change in methylome for *M. extorquens* populations growing with NaCl (200 mM, C) or with chloramphenicol (5 µg/ml) compared to without. **E)** Change in global DNA methylation pattern for different NaCl treatments (red = 0, light orange = 200, and dark orange = 250 mM). Figure shows best fit curves. Raw data shown in Fig S8. **F)** Change in global DNA methylation pattern for different chloramphenicol treatments (red = 0, light blue = 5, and dark blue = 10 µg/ml). Figure shows best fit curves. Raw data shown in Fig S9. **G)** Positive correlation between growth rate and strength of chromosome position dependent methylation pattern for *M. extorquens* (red, 𝑅^2^ = 0.65), *C. crescentus* (blue, 𝑅^2^ = 0.97) and *E. coli* (orange, 𝑅^2^ = 0.91). Green diamond represents *M. extorquens* with CcrM overexpressed (Fig 1H). Top inset - Illustration of the amplitude metric used for *M. extorquens* (Methods). Bottom inset – zoomed in on relationship for *E. coli*.

### Growth rate dependent shifts in global methylation patterns

To further test how patterns of DNA methylation depend on the environment, we characterized the methylome of *M. extorquens* populations growing in the presence of different stressors at sub-MIC levels. The methylation patterns in these environments were compared to that of growth in absence of the stressors. Stressful environments resulted in widespread shifts in methylation patterns with no clear outliers, in stark contrast to the minor, isolated changes observed on changing the substrate from succinate to methanol. For instance, salt stress significantly increased the average methylation state of most sites (Fig 2C), and chloramphenicol had similarly strong effects but in the opposite direction, as most sites exhibited significantly decreased methylation levels (Fig 2D).

By examining these shifts from the lens of chromosomal position, it became clear that nearly all of the effects of each stressor on methylation levels could be explained by their chromosomal position-dependent effects. Salt stress primarily increased average methylation levels around the replication origins (Fig 2E, Fig S8). In contrast, chloramphenicol led to widespread decreased methylation, especially strong close to the replication termini (Fig 2F, Fig S9). In both cases, increasing concentrations of stressor enhanced the change observed at the lower dose. Ciprofloxacin stress led to yet a distinct pattern, whereby some regions of the genome showed decreased methylation levels while others showed an increase (Fig S10).

Despite the distinct global changes in methylome for different environments, we observed that stressed cells tended to display a weaker chromosome position dependent methylation pattern (Figs 2E,F). In particular, there was a strong positive relationship between growth rate and the strength of the chromosome position dependent methylation pattern (Fig 2G, red), estimated as the distance between the minima and maxima for the best fit curves (Fig 2G top inset, Methods). To test the generality of these results, we studied the methylome of *C. crescentus* populations inhibited with varying concentrations of salt and chloramphenicol. Like for *M. extorquens*, these stressors had a global effect on methylation levels (Fig S11), and strength of the chromosome position dependent methylation pattern was strongly correlated with growth rate (Fig 2G, blue).

We postulated that the relationship between growth rate and position dependence of methylation patterns would be a unique feature of the cell-cycle dependent methylase activity seen in alphaproteobacteria. We examined the pattern of position dependence for GATC sites in *E. coli*, where expression of Dam methylase is constitutive, and observed that it was greatly diminished despite the wide range of growth rates (Fig 2G, orange and bottom inset). Constitutive CcrM expression in *M. extorquens* also reduced this pattern without much effect upon growth rate (Fig 1H, diamond in Fig 2G). All environments used in Fig 2G are listed in Table S5.

### Model of cell cycle can explain shifts in global methylation patterns

We postulated that the effects of different environments on global methylation patterns can be shaped simply by their distinct effects on the cell cycle. We developed a simple quantitative model of cell division to try to predict the expected changes in methylation along the chromosome for stressors that impact distinct aspects of the cell cycle (Fig 3A). Cell division for bacteria is typically divided into three stages^43^ – B (initiation), C (elongation), and D (division), corresponding to the period before, during, and after DNA replication. During B phase, we assume all methylation sites to be fully methylated. During C phase i.e., during DNA replication, the proportion of time each locus spends in a fully-methylated vs hemi-methylated state is a function of the distance from the origin of replication. CcrM is transiently active between late C and early D phases, and is known to methylate one site at a time^44^. We can then define an average waiting time for each site before it is fully-methylated again. Based on this framework, we arrive at analytical equations (Methods) to estimate the fraction of time each site is methylated based on position in the chromosome (Fig 3B). The model can capture the observed chromosome position dependent pattern of DNA methylation (Fig 3C).

**Fig 3.**
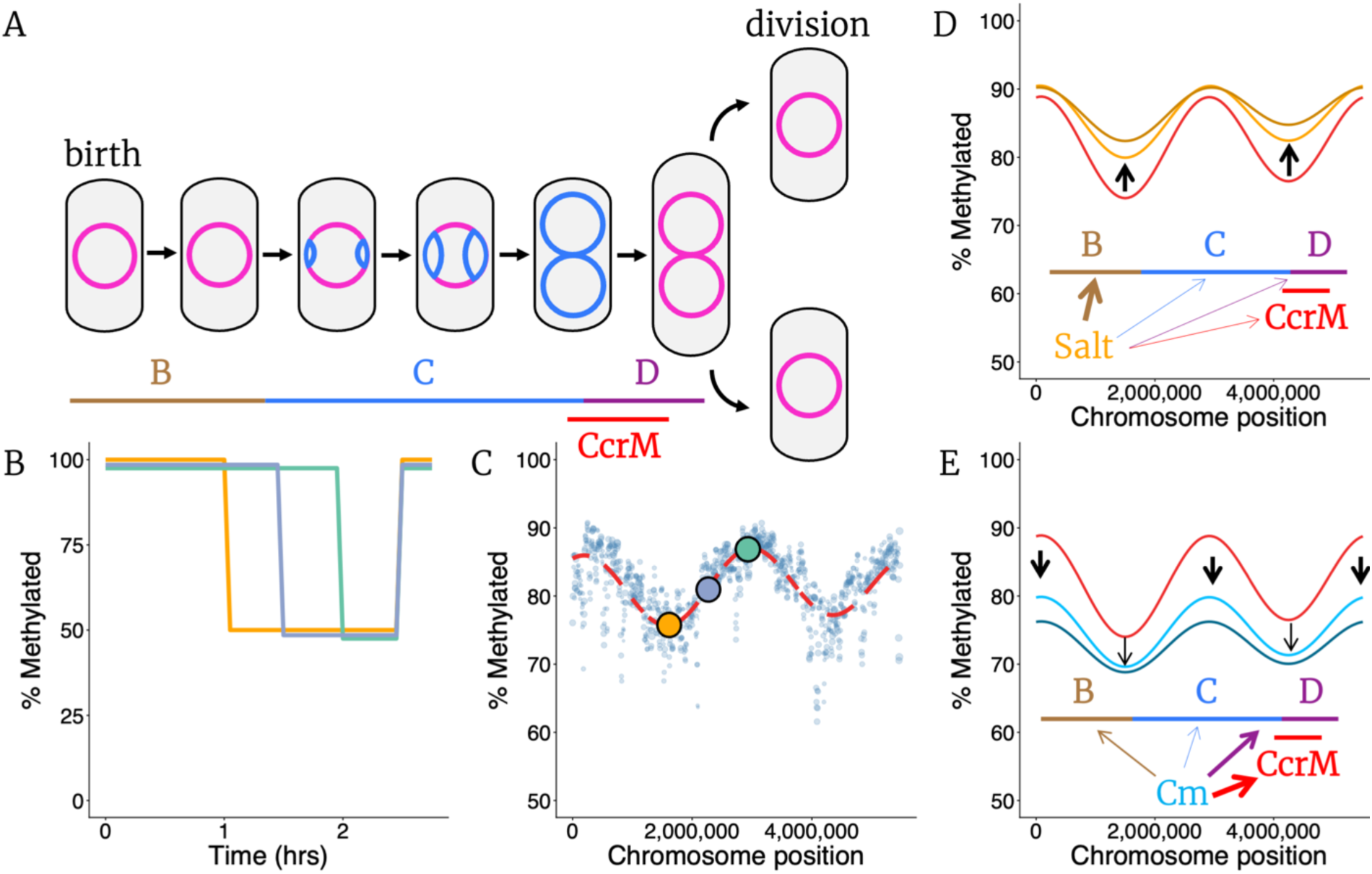
Model of cell cycle can explain the response of global methylation levels to environmental shifts. **A)** Cell cycle is divided into three phases – B, C, and D, corresponding to the periods before, during, and after DNA replication. The three phases are shown using solid bars below the cells, and new hemi-methylated strands of DNA shown in blue. CcrM is transiently active towards the beginning of the D phase depicted by the red bar during which new strands are remethylated. **B)** Theorized average methylation states of sites in different genomic regions as a function of the cell cycle (green – close to terminus, orange – close to origin, blue – midway between the two). The lines are offset for visibility. All sites start the cell-cycle fully methylated, become hemi-methylated after passage of the replication fork and are re-methylated some period after CcrM is active. However, the methylation dynamics are distinct as a function of distance from the origin of replication. **C)** The predicted pattern of methylation as a function of genomic position based on our model (parameters used are listed in Table S6). Dashed red line represents best fit curve based on model predictions, and points represent empirical data during growth on succinate. Colored points represent the positions used for panel B. **D)** Theoretical predictions of chromosome position dependent methylation patterns generated for populations predominantly inhibited in the B phase (thickness of arrows for various aspects of cell cycle represent relative impact on those phases) are similar to patterns seen for salt stress for *M. extorquens*. Darker colors represent increased inhibition. Red curve corresponds to prediction for no inhibition. **E)** Theoretical predictions of chromosome position dependent methylation patterns generated for populations with inhibited CcrM activity and longer D phase (thickness of arrows for various aspects of cell cycle represent relative impact on those phases) are similar to patterns seen for chloramphenicol (Cm) stress. Darker colors represent increased inhibition. Red curve corresponds to prediction for no inhibition.

Inhibiting each of the phases of the cell cycle is predicted to lead to a different canonical change in the pattern of methylation along the chromosome, thereby serving as a potential diagnostic indicator of which stressors most impact which aspects of physiology (Methods, Fig S12). For instance, increases in time spent in B and/or D phases leads to a global increase in methylation level, which is consistent with the pattern observed for salt stress (Fig 3D). As another example, the observed patterns of decreased methylation levels seen for chloramphenicol can be explained by an increase in B/D phases coupled with reduced CcrM activity (Fig 3E). This results in an increase in the waiting time for sites in the new strand to be methylated relative to the length of D phase. That chloramphenicol generates an opposite response as salt is perhaps unsurprising, given that recent experiments in *E. coli* have shown that cellular response to slow growth depends on the mechanism of growth inhibition and is particularly distinct for translation inhibition^45,46^. We can similarly explain the effect of ciprofloxacin on methylation patterns based on the known effect on DNA damage and inhibiting cell division (Fig S13).

## Discussion

The growth rate dependence of methylation patterns observed across organisms illustrates the possibility of using methylome sequencing data as a tool to learn about microbial physiology in natural microbiomes. In addition to estimating growth rate, the patterns of methylation across the genome can also serve as an indicator about which cellular processes are most impacted in which environment. For instance, the effect of salt stress on *Methylobacterium* (Fig 2E) is explained by inhibition in B/D phases (Fig 3D), similar to previous work that slow growth, albeit caused by nutrient limitation, has a generic impact on increasing time needed for all stages of cell division but predominantly the B phase^47^. The effect of chloramphenicol stress (Fig 2F) is explained by an increase in average waiting time for a site to be methylated (Fig 3E). A decrease in CcrM activity suggested by this pattern could be due to changing levels of global transcriptional regulators in response to chloramphenicol^48,49^ which either act on CcrM directly, or change CcrM substrate pools like *S*-adenosyl methionine (SAM). These ideas complement existing approaches on estimating growth rate based on differences in sequencing coverage across the genome^25,26^. However, methylome data provides a more robust estimate for slow-growing microbes where the differences in coverage between origin and terminus are minimal. Additionally, methylation patterns can also provide information about the mode of growth inhibition in natural settings, with different classes of slow growth exhibiting distinct methylation patterns.

Our results also highlight the presence of multiple origins of replication in *M. extorquens*. This is surprising since bacterial genomes are widely thought to possess a single origin. However, one strain of *Agrobacterium tumefaciens* has been shown to switch between having two chromosomes and a single fused chromosome with multiple origins^50^. Similar patterns of stable chromosome fusion and multiple origins of replication were recently observed in clinical isolates of *Vibrio cholerae*^38^. Together with our observations, these results suggest the presence of multiple origins of replication in bacteria might be much more common than previously thought, and future work is needed to understand the prevalence, origin, and maintenance of such genomic rearrangements.

For GANTC motifs in alphaproteobacteria, all genes previously shown to be under epigenetic regulation have been cell-cycle regulated where gene expression changes as methylation sites transition from fully methylated to hemi-methylated as DNA replication progresses. The switch in methylation pattern observed for sites upstream of formate dehydrogenase when switching between succinate and methanol hints at a first example of phase variation associated with GANTC motifs. Such patterns of pairs of methylated motifs in a promoter sequence switching together have been frequently seen in phase-variable genes associated with GATC motifs^51^. An understanding of the gene expression dynamics and transcription factors involved will help understand the role and significance of such switches in alphaproteobacteria and in particular, implications for methylotrophic metabolism.

A chromosome position dependent DNA methylation pattern has been consistently observed for GANTC sites in alphaproteobacteria, and dozens of genes have been shown to significantly change expression levels as the proportion of time spent in fully methylated vs hemi-methylated states change, as tested by synthetically modifying either CcrM levels^13^ or the distance of a gene from replication origin^11^. Here, we find that different growth regimes can naturally change these position dependent patterns, leading to significant undermethylation like with chloramphenicol, or close to fully-methylated for salt stress. Both undermethylated and fully-methylated phenotypes have been previously characterized for several alphaproteobacteria using genetic constructs, finding significant changes in expression levels of hundreds of genes. The most consistent observations have been reduced methylation changing expression of genes essential for cell division and increased methylation altering expression of stress response and metabolic regulons. Our results show that growth inhibition alone, via distinct physiological effects, can change global methylation patterns in the direction of either significantly reduced or increased methylation levels in certain regions of the genome, potentially modifying gene expression to combat the mode of inhibition. These results also suggest DNA methylation as a novel force shaping genome architecture with the location of a gene in the genome differently impacting gene expression for different growth conditions.

Studying the methylome for different organisms across different environments, we find that methylation patterns across the genome are highly fluid with average methylation states changing significantly for a large number of loci. However, it remains unknown how rapidly these global changes revert once the environmental stressor is removed. It was recently discovered that *P. aeruginosa* exhibits global hypomethylation upon increased temperature, and the global patterns persist for dozens of generations after temperature has decreased^53^. There is also recent evidence that even in constant environments, methylation states are not static and can rapidly evolve over short timescales^54,55^. Further work is needed to characterize both the immediate and long-term evolution of methylome patterns in response to diverse environments. Environmental dependence of methylation patterns may also be prominent in eukaryotic genomes, where similar principles of cell division are known to shape global epigenetic patterns^56^.

## Methods

### Bacterial culturing and sequencing

The WT *Methylobacterium* strain used in this work is a mutant of *Methylobacterium extorquens* PA1^57^, deleted for cellulose biosynthesis genes (Δ*cel*), which aids growth in laboratory conditions. All cultures were grown in *Methylobacterium* PIPES (MPIPES) minimal media^58^, supplemented with either succinate (15 mM) or methanol (30 mM) as a carbon source. All stressors were tested in succinate-grown cultures. *C. crescentus* NA1000 cultures were grown on peptone yeast extract (PYE, Table S7). *E. coli* MG1655 cultures were grown on M9 minimal media with 4 mM glucose as the carbon source. All cultures were grown in test tubes in volumes of 5 ml, shaking at 250 rpm, at 30 °C for *M. extorquens* and *C. crescentus*, and 37 °C for *E. coli*. For constitutive *ccrM* expression, the gene (Mext_3181) was cloned into a plasmid and regulated using a *tetR* repression system described previously^59^. To characterize the effect of constitutive CcrM activity, growing cells were induced using 50 nM anhydrous tetracycline (aTc) and cells harvested 90 min after induction to quantify the methylome (Fig 1H).

For sequencing the methylome, exponentially growing cells were centrifuged at 4000g for 10 min, washed with PBS, centrifuged again at 4000g for 10 min, and the pellet resuspended in 500 µl DNA/RNA Shield (Zymo Research, USA). DNA extraction and long-read nanopore sequencing was performed by Plasmidsaurus using R10.4.1 flow cells and raw POD5 data delivered.

### Methylome analysis

Dorado super-accurate model v5.2.0^60^ was used for basecalling using a custom bash script, outputting modified bam files associated with each sample. We used minimap2^61^ to align reads to the reference genome and modkit^62^ to estimate the number of reads in which each nucleotide was predicted to be methylated. The output of the modkit data was processed using custom scripts in R to analyze the fraction of times a particular site was methylated or unmethylated.

### Discovery of methylation motifs

We used the protocol outlined in Tidwell et al^63^ to identify methylated motifs from our sequencing data. In brief, bedmethyl files produced using modkit were analyzed using MEME^64^ to detect genetic contexts in which a nucleotide is consistently methylated. The same methylation motifs were also independently identified using the motif search function in modkit.

### Cumulative GC skew calculations

Cumulative GC skew^34,35^ is defined as the bias in presence of G over C on one strand of the DNA. For every window of 5000 bp, we estimate this bias 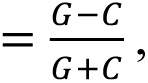 where G and C simply represent the sum of all guanines and cytosines in that stretch of DNA. A rolling sum of this bias over the whole genome is represented as cumulative GC skew (Fig 1, Fig S3). This approach helps identify the replication origin and terminus as the leading strand during DNA synthesis is observed to almost universally display a bias towards G over C.

### Statistical Analysis

For comparing differences in methylation states between samples, Fisher’s exact test was coupled with Benjamini-Hochberg procedure to correct for false discovery rate and calculate *p*-values associated with each site for each pair of samples. A threshold of *p* < 0.05 was used to classify sites as statistically significant. For Fourier regression, the methylation vs chromosome position data (Fig 1D) was fit using the following 2^nd^ order function: 𝑦 = 𝐴_0_ + 𝐴_1_. cos(𝑥) + 𝐵_1_. sin(𝑥) + 𝐴_2_. (𝑐𝑜𝑠2𝑥) + 𝐵_2_. (𝑠𝑖𝑛2𝑥). The function was limited to just two harmonics because of the visual periodicity of the data with two peaks and troughs, and the underlying biology of two replication origins and termini. The best fit coefficients were estimated using linear regression and the associated curve shown in red in Fig 1D. The same analysis was applied to data from other environments (Fig 3). The strength of the chromosome position dependent pattern of the experimental data was estimated using the difference in methylation between the origins and termini (average methylation at termini for *M. extorquens*), whose locations were calculated based on the first derivative of the best fit function being zero. For *C. crescentus*, the methylation vs chromosome position data (Fig 1A, Fig S11) was fit using the single-order function: 𝑦 = 𝐴_0_ + 𝐴_1_. cos(𝑥) + 𝐵_1_. sin(𝑥). All statistical tests were performed in R (version 4.5.2).

### Model of cell division

Cell division is divided into three phases (Fig 3A). Consider any one methylation site on the genome. During B phase, the site remains full-methylated. During C phase, the site becomes hemi-methylated after passage of the replication fork. The time taken for the site to become hemi-methylated is thus a function of its position on the genome. After passage of the replication fork, the site remains hemi-methylated until CcrM is active and remethylates the site. Using this framework, we can define the average methylation state for a site using the following equation –

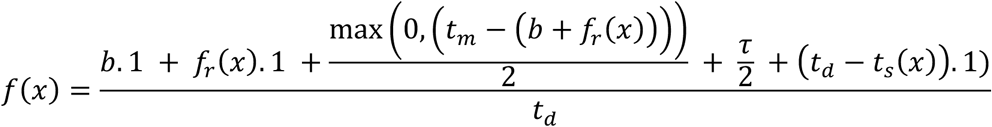

where *x* represents the normalized position of the site on a replication arm (between 0 and 1, with 0 being the origin and 1 being the terminus). *b* represents the time spent in the B phase. 𝑓_r_(𝑥) denotes the time after the start of DNA replication for which the site remains full-methylated. In our model, 𝑓_r_(𝑥) = 𝑐. 𝑥, with *c* representing the time taken for DNA replication on that arm. 𝑡_m_ represents the time after the start of the cell cycle before the methylase (CcrM) is active.

τ represents the average waiting time for a site before methylase activity, 𝑡_d_ denotes doubling time, and 𝑡_s_ denotes time taken for all processes described so far.

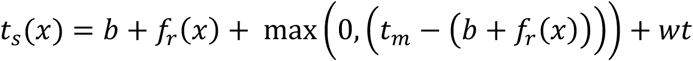

For the second origin, replication is assumed to begin some time *α* after the first, based on observations in *Agrobacterium tumefaciens*, a related alphaproteobacteria with two origins^50,65^. Sites linked with the second origin of replication thus spend more time in the ‘B’ phase and we can simply use: 𝑏, = 𝑏 +. These equations were used to estimate average methylation for many locations on the genome and the best fit curve estimated. This approach predicts a similar pattern to the chromosome position dependent pattern observed empirically (Fig 3C). Parameters used for simulations are listed in Table S6.

To understand the role of each of these variables in shaping the methylome, we tested perturbations to these parameters, and quantified the change in methylome profile. Different perturbations had distinct and unique effects on the pattern of methylation (Fig S12), and understanding the effect each variable had on the methylation pattern helped formulate the relative effect of environmental change on each variable. For salt stress, we postulated a small increase in all steps of the cell cycle but a major increase in the B phase, *b* (Fig 3D). For chloramphenicol, we postulated major increases in the D phase *d*, and the average waiting time, τ for a site to be methylated (Fig 3E).

## Acknowledgements

We are grateful to Olin Silander, Megan Behringer, and Adam Guss for help with analysing nanopore sequencing data. We thank Kevin Gozzi for sharing the *Caulobacter crescentus* NA1000 strain. We thank members of the Marx and Udekwu labs for discussions and helpful feedback on the manuscript. We thank Deena B. Goodgold for help with making figures.

## Funding

This work was funded by Department of Energy (DOE) awards DE-SC0026232 to KIU and CJM, and DE-SC0022318 to CJM.

## Data Availability

Sequencing data will be deposited on NCBI SRA upon publication. All scripts used in this work are available on GitHub – https://github.com/mallakshat/Methylome_GrowthRate

## Supplementary Information

### Supplementary Methods

#### Estimations of bias in distribution of methylation motifs

For estimating bias in presence of methylation motifs in the genome (Fig S1), we first estimated the expected number of sites for each motif in the genome based on a random distribution of nucleotides over the length of the genome. For example, in a random DNA sequence, any nucleotide can be the start of a GANTC site with a probability = 1/(4^4^). This probability was corrected for GC bias for all strains analyzed. Multiplying the probability by the length of the coding/intergenic regions tells us the expected number of GANTC sites. Bias was quantified as the ratio of observed number of methylation sites to this expected number.

---

**Fig S1.**
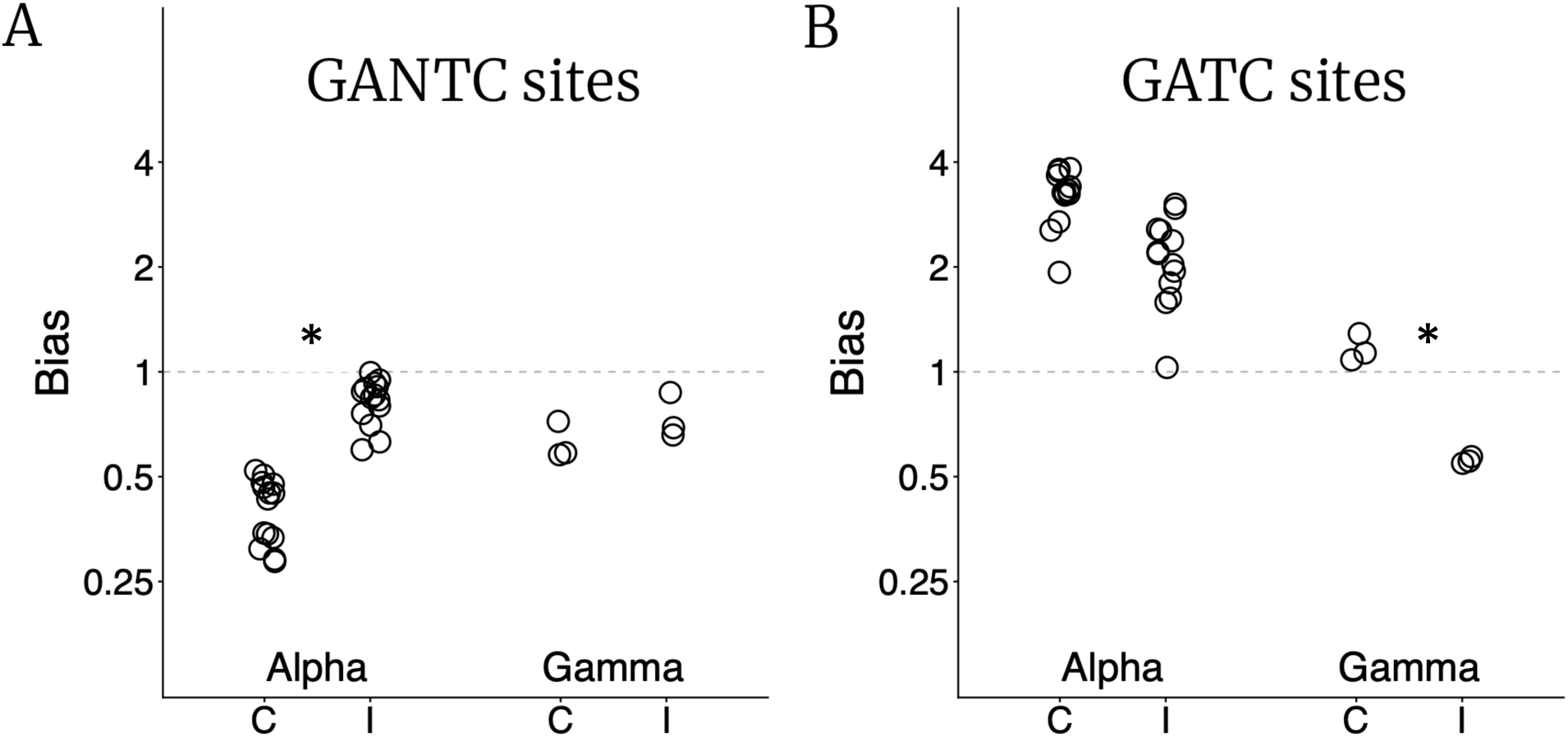
Bias in distribution of methylation motifs. Biases are defined as the ratio of observed number of sites for a motif relative to the expected number (Supplementary Methods). Plots shown on a log_2_ scale (y-axis). **A)** Bias in presence in coding (C) and intergenic (I) regions for GANTC sites (targets of methylation in alphaproteobacteria) in alphaproteobacteria and gammaproteobacteria. **B)** Bias in presence in coding (C) and intergenic (I) regions for GATC sites (targets of methylation in gammaproteobacteria) in alphaproteobacteria and gammaproteobacteria.

**Fig S2.**
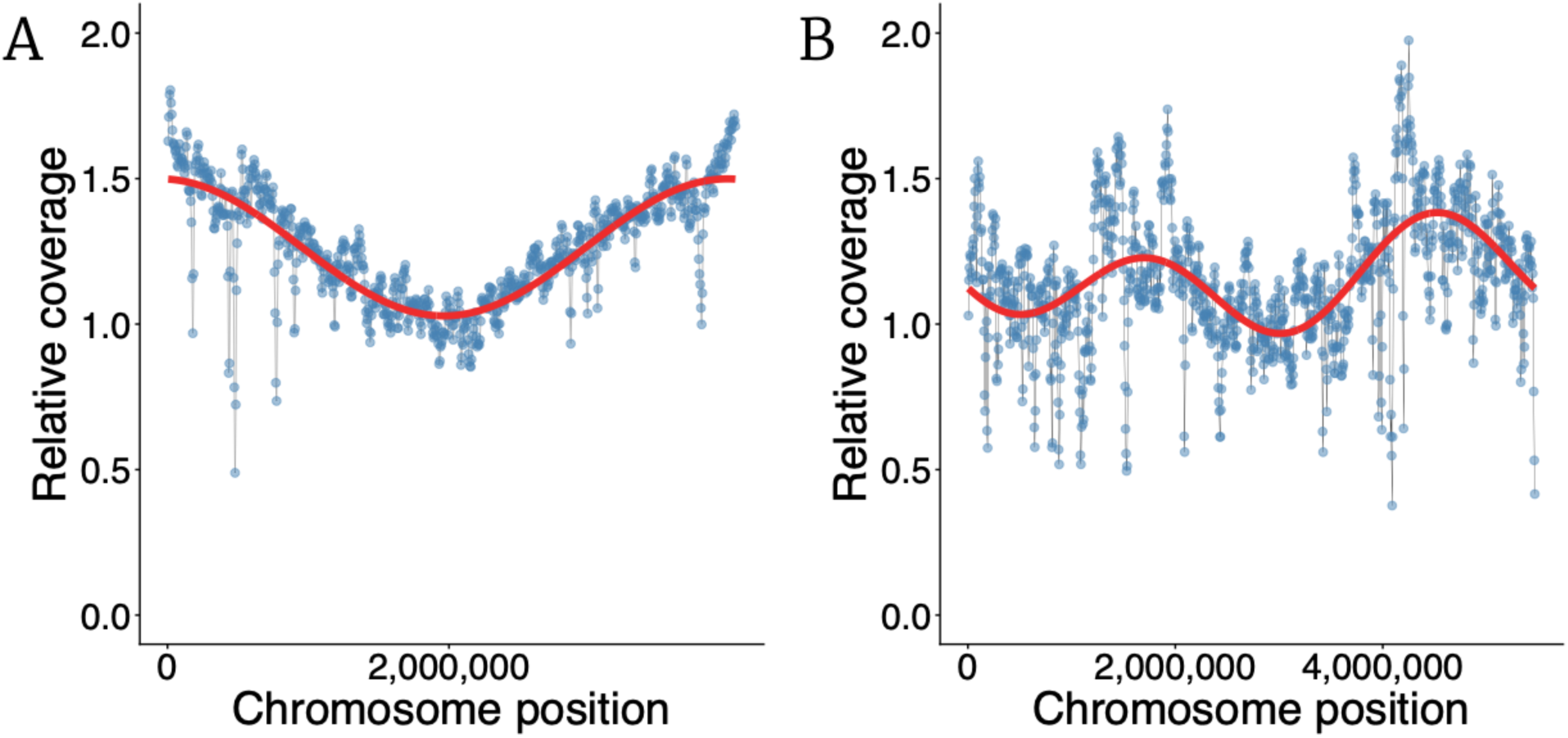
Relative coverage vs position for. **A)** *Caulobacter crescentus* and **B)** *Methylobacterium extorquens*. Relative coverage is defined as the number of reads for a given position relative to the number of reads around the replication terminus. Each point represents the average relative coverage in a 10,000 bp window around that point (points separated by 5000 bp). Red curves denote best fit from a Fourier regression (Methods). Sequencing coverage shows the same pattern of replication origins and termini as methylation patterns (Fig 1E,F).

**Fig S3.**
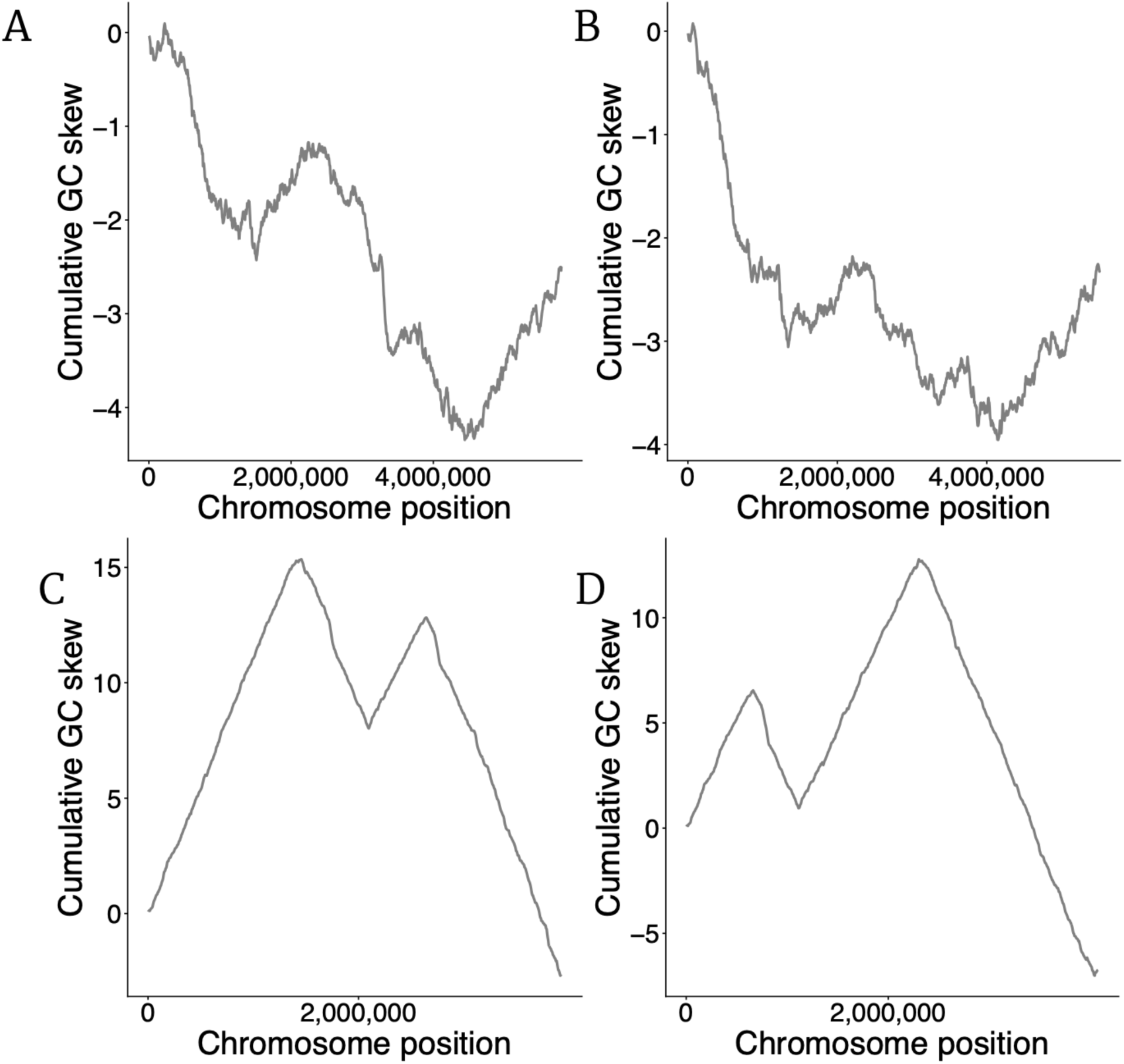
Cumulative GC skew for. **A)** *Methylobacterium populi* BJ001. **B)** *Methylobacterium extorquens* AM1. **C)** *Vibrio cholerae* 1154-74. **D)** *Vibrio cholerae* 10432-62. Some *Methylobacterium* strains (panels A,B) show a similar pattern, like *M. extorquens* PA1 (Fig 1F), of multiple origins of replication. *Vibrio cholerae* strains known to possess two origins show a similar pattern (panels C,D). Cumulative GC skew calculations are explained in the Methods.

**Fig S4.**
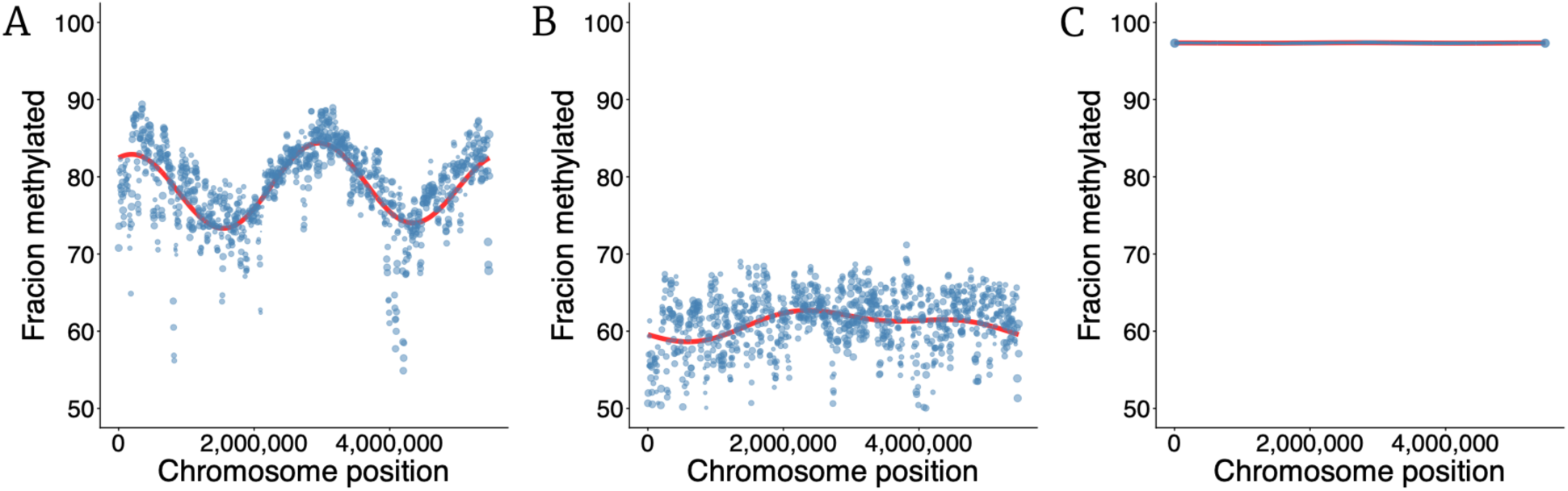
Fraction methylation vs chromosome position. Chromosome position dependent pattern of fraction of methylated sites. Each point represents average proportion of methylated reads for all sites in a 10,000 bp window around that point (points separated by 5000 bp). Size of points are proportional to the number of sites in that window. Red curves denote best fit from a Fourier regression (Methods). **A)** GANTC sites for cells growing exponentially on methanol. **B)** GANTC sites for cells in stationary phase after growth on succinate. **C)** GCCGATC sites for cells growing exponentially on succinate.

**Fig S5.**
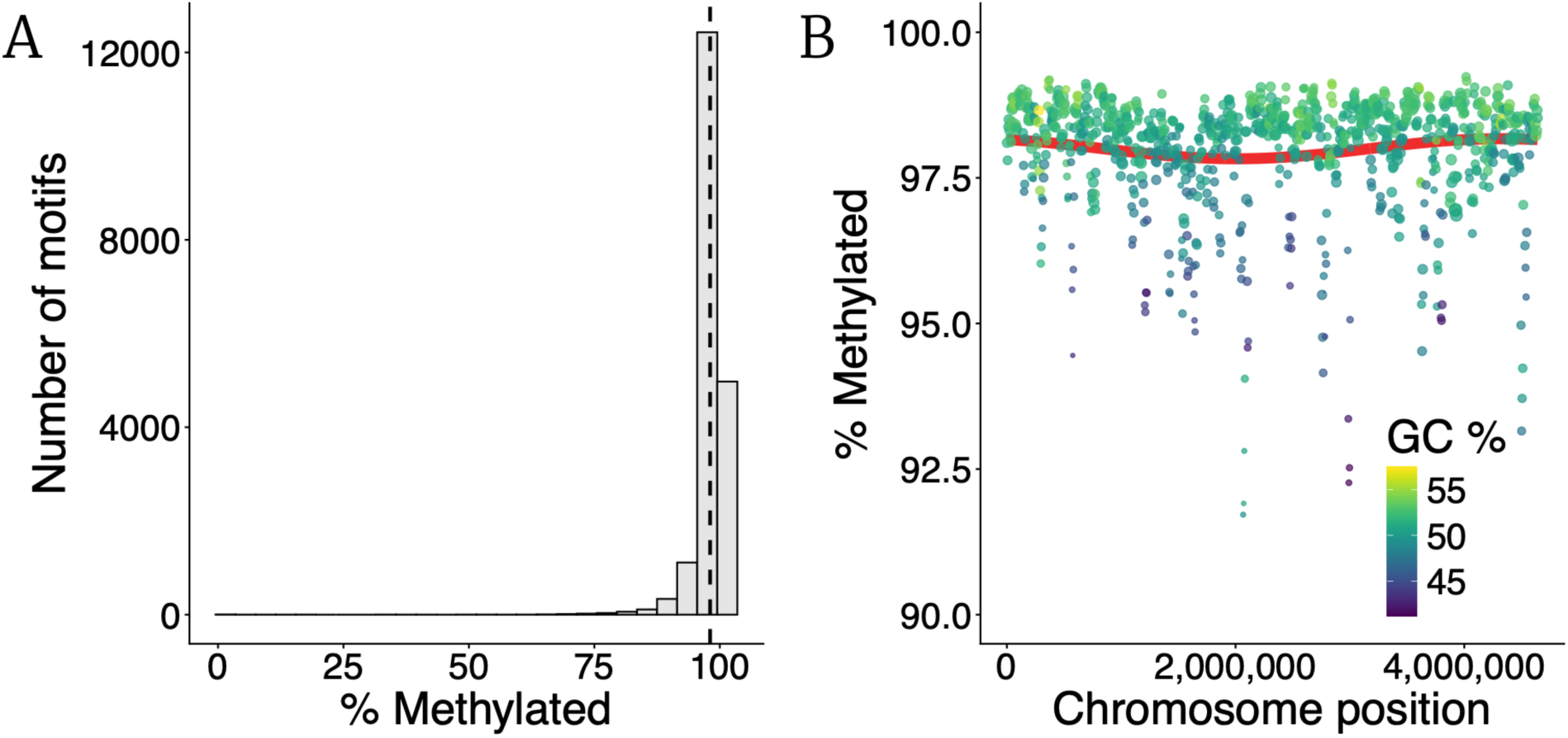
Methylation patterns for GATC sites in *E. coli*. **A)** Distribution of number of sites which are seen methylated a particular fraction of times. **B)** Chromosome position dependent pattern of methylation. Each point represents average proportion of methylated reads for all sites in a 10,000 bp window around that point (points separated by 5000 bp). Size of points are proportional to the number of sites in that window. Color of the points represent average GC content in the 10,000 bp window around that point. Red curve denote best fit from a Fourier regression (Methods).

**Fig S6.**
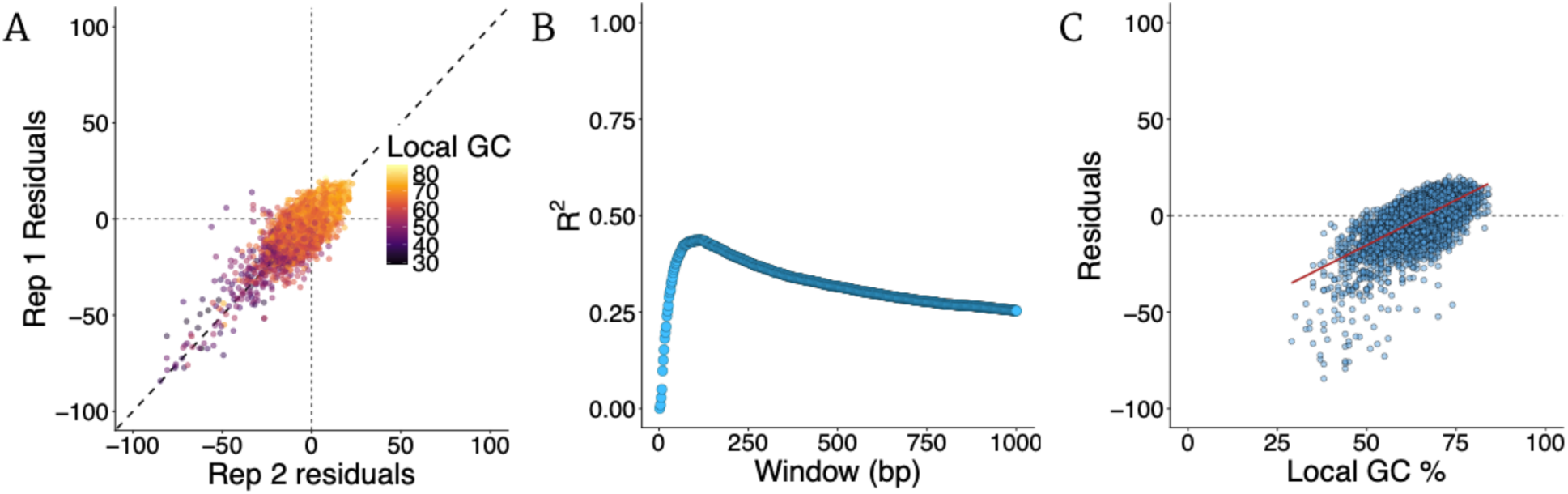
Local GC percent explains variance around position dependent methylation pattern for *M. extorquens*. **A)** Scatter plot of residuals from chromosome position dependent pattern (Fig 1C) between replicate populations growing exponentially on succinate. Each point represents one GANTC site. Dashed line represents the x=y line. Residuals for a site are defined as distance of average methylation state from the expected (fitted red curve in Fig 1C). Points are colored based on GC content in a 100 bp window around that site. **B)** Coefficient of determination (𝑅^2^) for proportion of variance explained by local GC content as a function of the number of neighboring nucleotides considered around a site. ∼100 bp window around each site explains maximum variance in the pattern of residuals vs local GC content. **C)** Scatter plot of residuals vs local GC for a window of size 100 bp. Best linear fit denoted in red.

**Fig S7.**
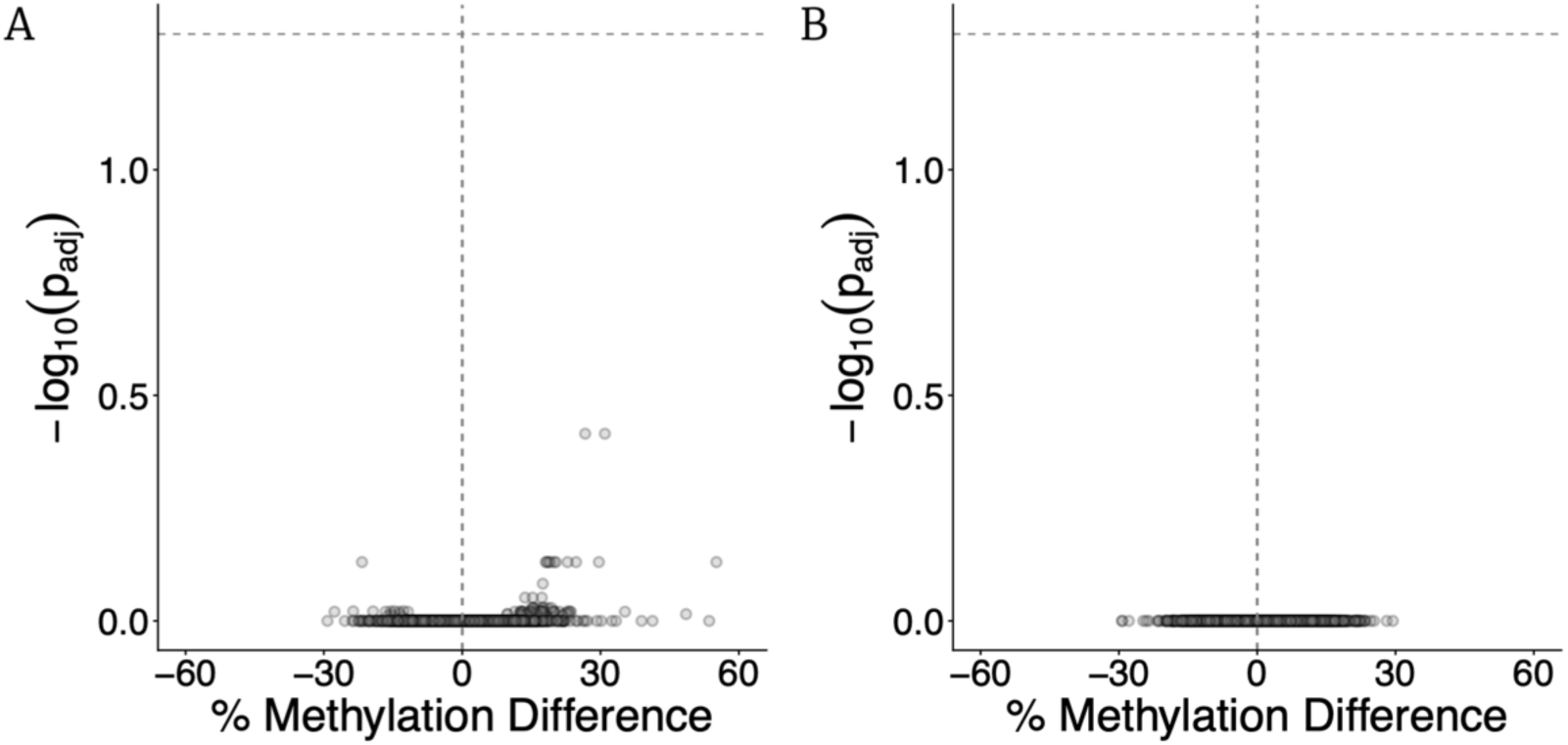
No change in methylome between replicate populations in the same environment. **A)** Change in methylation for replicate populations growing on succinate. Each point represents the observed change in fraction of methylated reads (x-axis) for one GANTC site and the *p*-value associated with the change (y-axis). **B)** Change in methylation for replicate populations growing on methanol. Each point represents the observed change in fraction of methylated reads (x-axis) for one GANTC site and the *p*-value associated with the change (y-axis).There are no statistically significant differences in methylation between replicate populations growing on either substrate.

**Fig S8.**
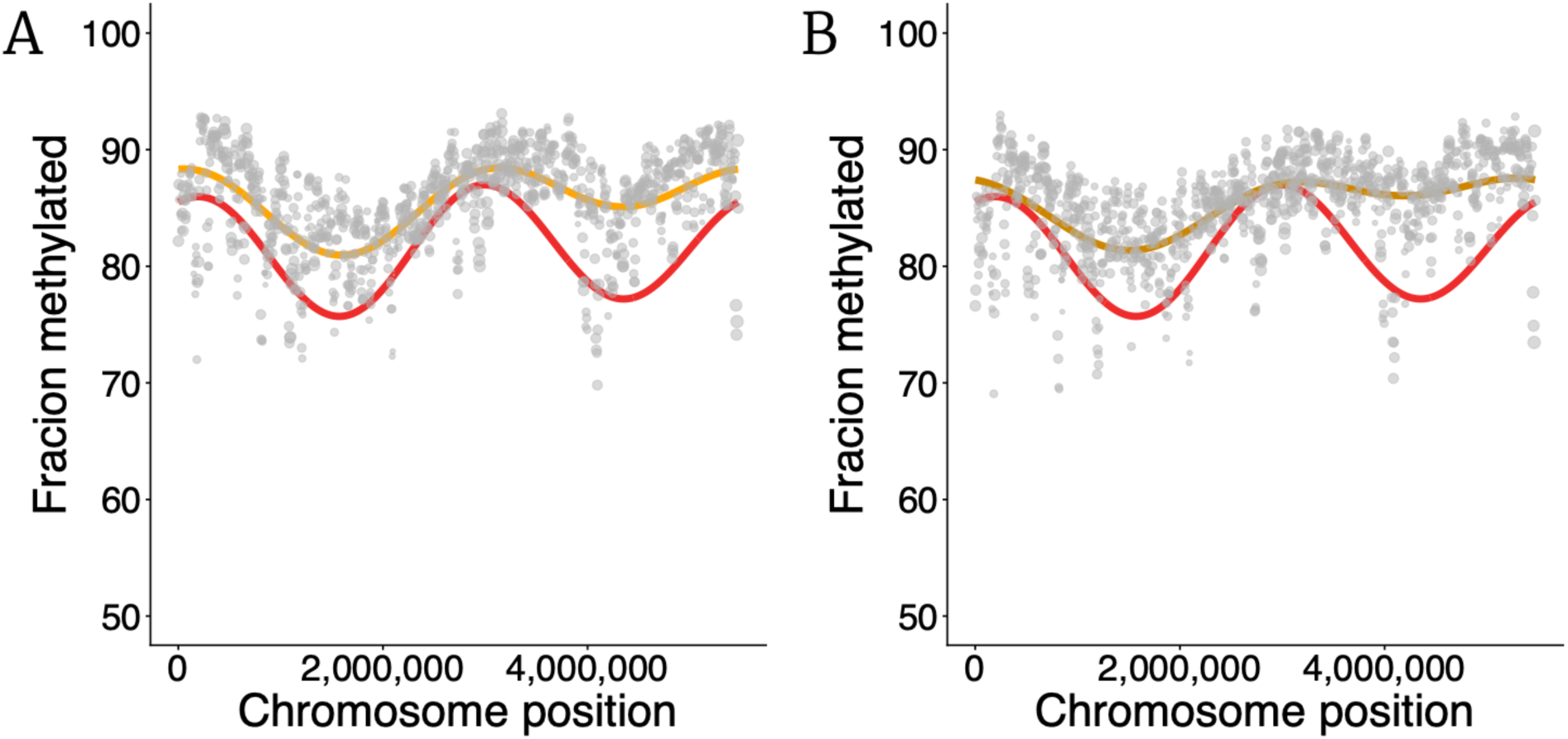
Chromosome position dependent pattern of fraction of methylated GANTC sites during growth with salt stress. **A)** 200 mM NaCl. **B)** 250 mM NaCl. Each point represents average proportion of methylated reads for all GANTC sites in a 10,000 bp window around that point (points separated by 5000 bp). Size of points are proportional to the number of GANTC sites in that window. Red curve shows best fit from a Fourier regression for growth on just succinate (same as Fig 1C). Orange curves show best fit best fit from a Fourier regression for each concentration of salt.

**Fig S9.**
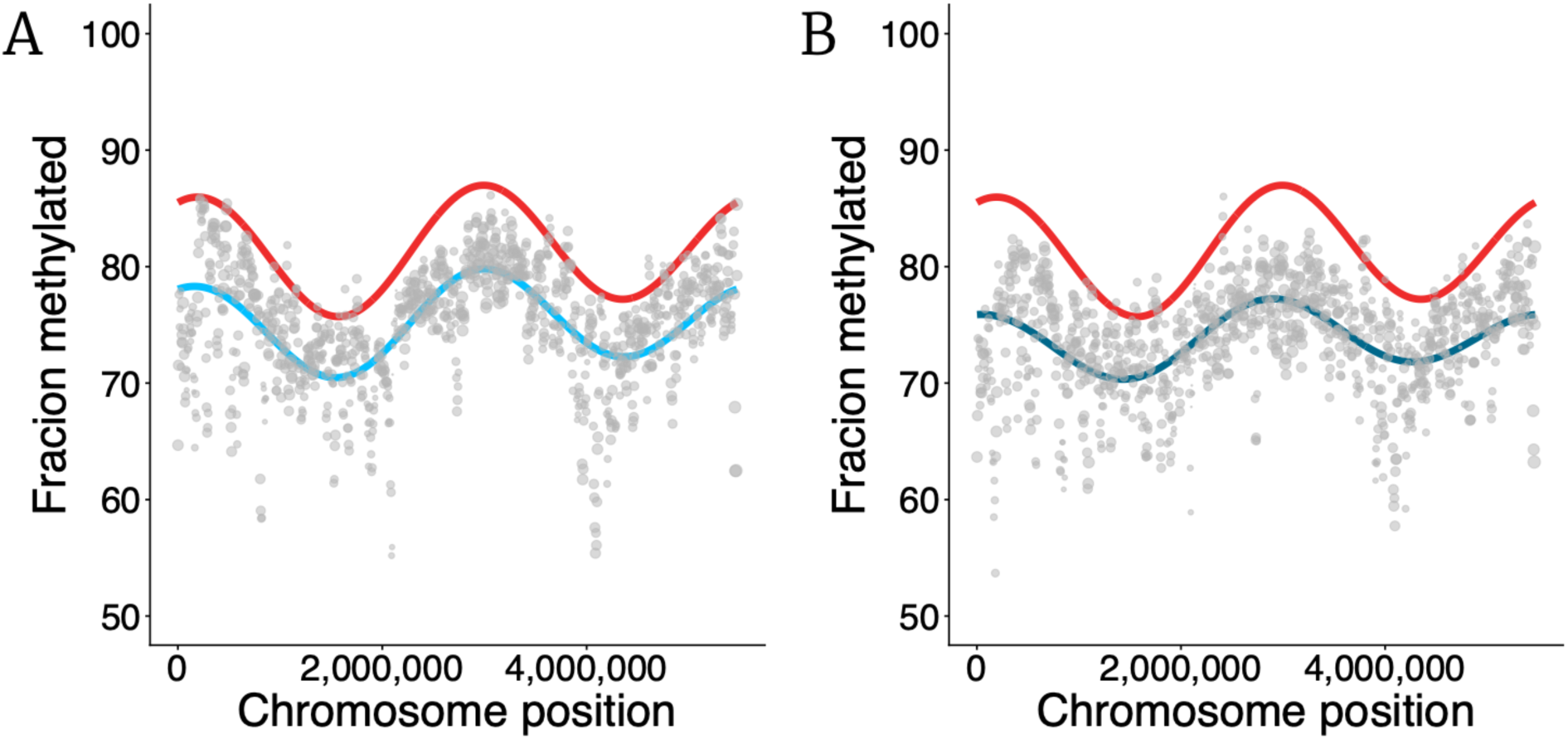
Chromosome position dependent pattern of fraction of methylated GANTC sites during growth with chloramphenicol. **A)** 5 µg/ml chloramphenicol. **B)** 10 µg/ml chloramphenicol. Each point represents average proportion of methylated reads for all GANTC sites in a 10,000 bp window around that point (points separated by 5000 bp). Size of points are proportional to the number of GANTC sites in that window. Red curve shows best fit from a Fourier regression for growth on just succinate (same as Fig 1C). Blue curves show best fit best fit from a Fourier regression for each concentration of chloramphenicol.

**Fig S10.**
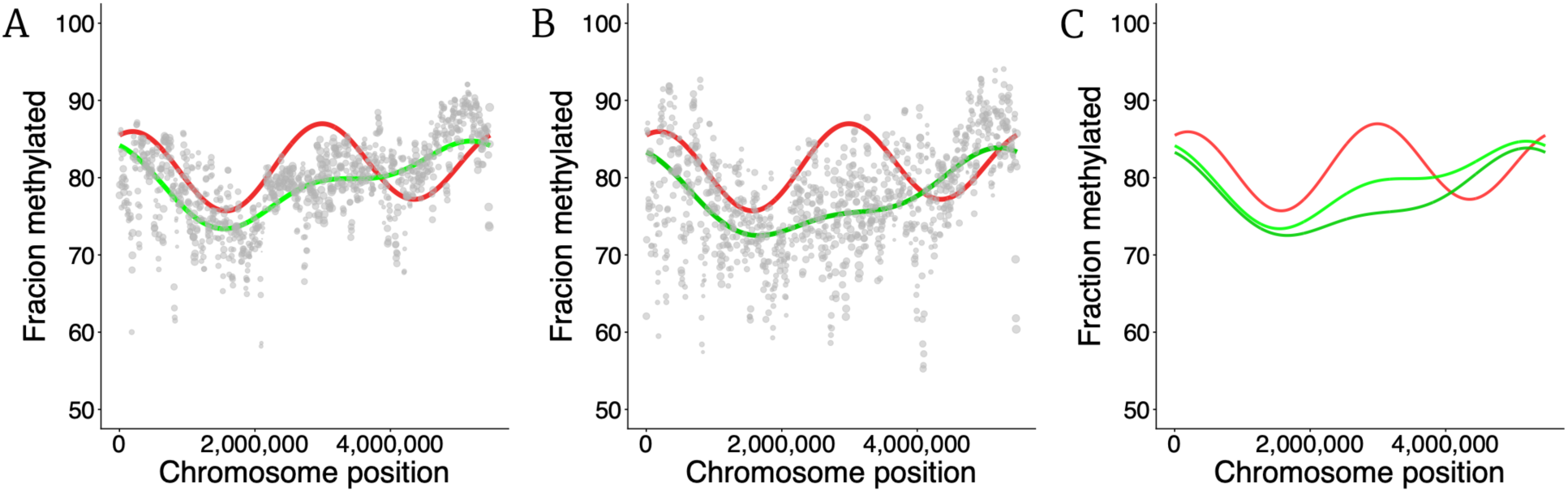
Chromosome position dependent pattern of fraction of methylated GANTC sites during growth with ciprofloxacin. **A)** 2 µg/ml ciprofloxacin. **B)** 4 µg/ml ciprofloxacin. **C)** Showing both concentrations together. In panels A and B, each point represents average proportion of methylated reads for all GANTC sites in a 10,000 bp window around that point (points separated by 5000 bp). Size of points are proportional to the number of GANTC sites in that window. Red curve shows best fit from a Fourier regression for growth on just succinate (same as Fig 1C). Green curves show best fit best fit from a Fourier regression for each concentration of ciprofloxacin. In panel C, the different ciprofloxacin treatments are shown together (red = 0, light green = 2, and dark green = 4 µg/ml).

**Fig S11.**
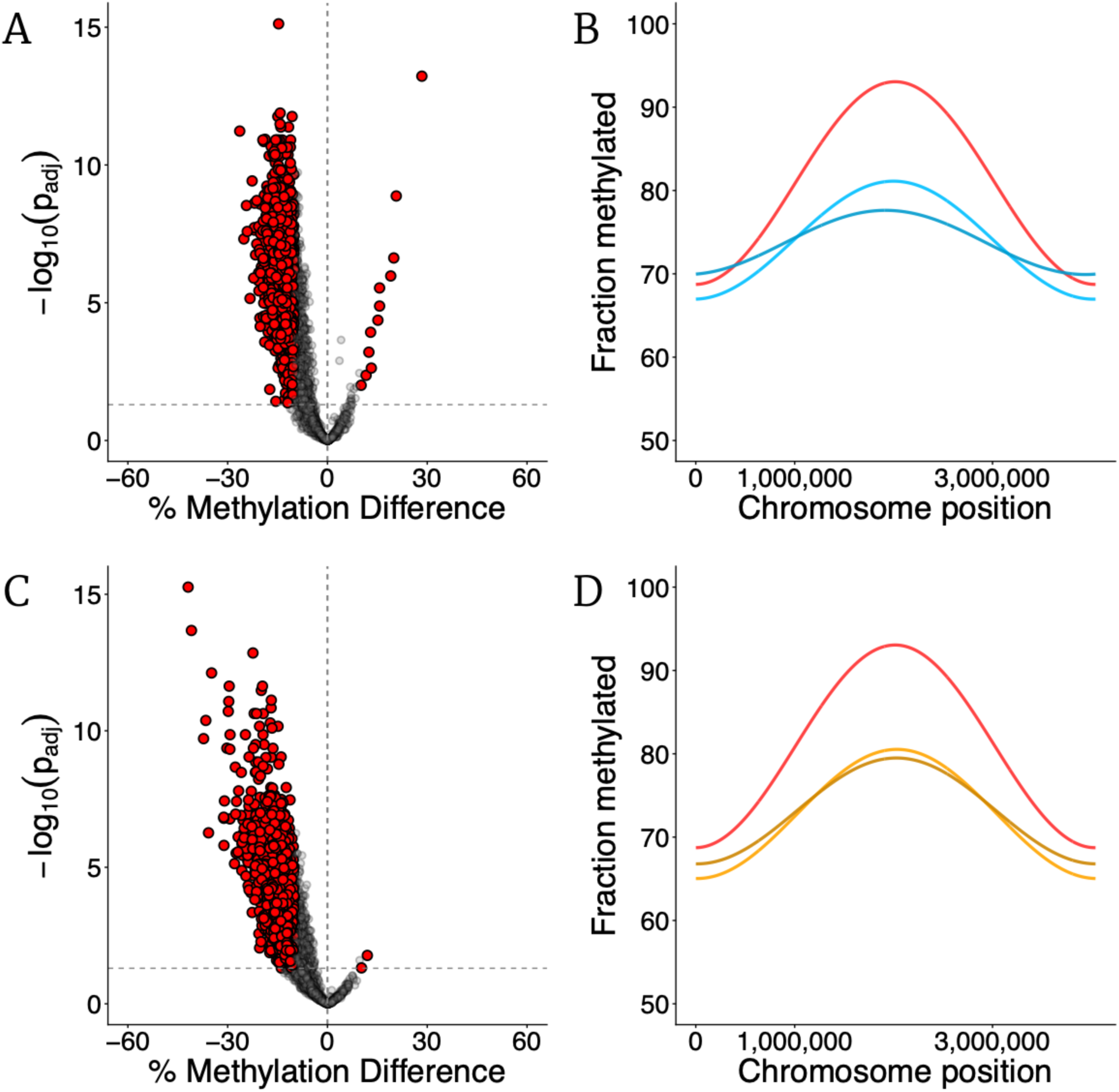
Change in *C. crescentus* global methylation patterns in different environments. **A)** Volcano plot comparing the change in methylation state for all GANTC sites during growth with 0.3 µg/ml chloramphenicol compared to without. Each point represents the observed change in fraction of methylated reads (x-axis) for one GANTC site and the *p*-value associated with the change (y-axis). **B)** Change in global DNA methylation pattern for different chloramphenicol treatments (red = 0, light blue = 0.3, and dark blue = 0.5 µg/ml). Data represented shows best fit curves estimated using Fourier regression. **C)** Volcano plot comparing the change in methylation state for all GANTC sites during growth with 90 mM NaCl compared to without. Each point represents the observed change in fraction of methylated reads (x-axis) for one GANTC site and the *p*-value associated with the change (y-axis). **D)** Change in global DNA methylation pattern for different NaCl treatments (red = 0, light orange = 90, and dark orange = 95 mM). Data represented shows best fit curves estimated using Fourier regression.

**Fig S12.**
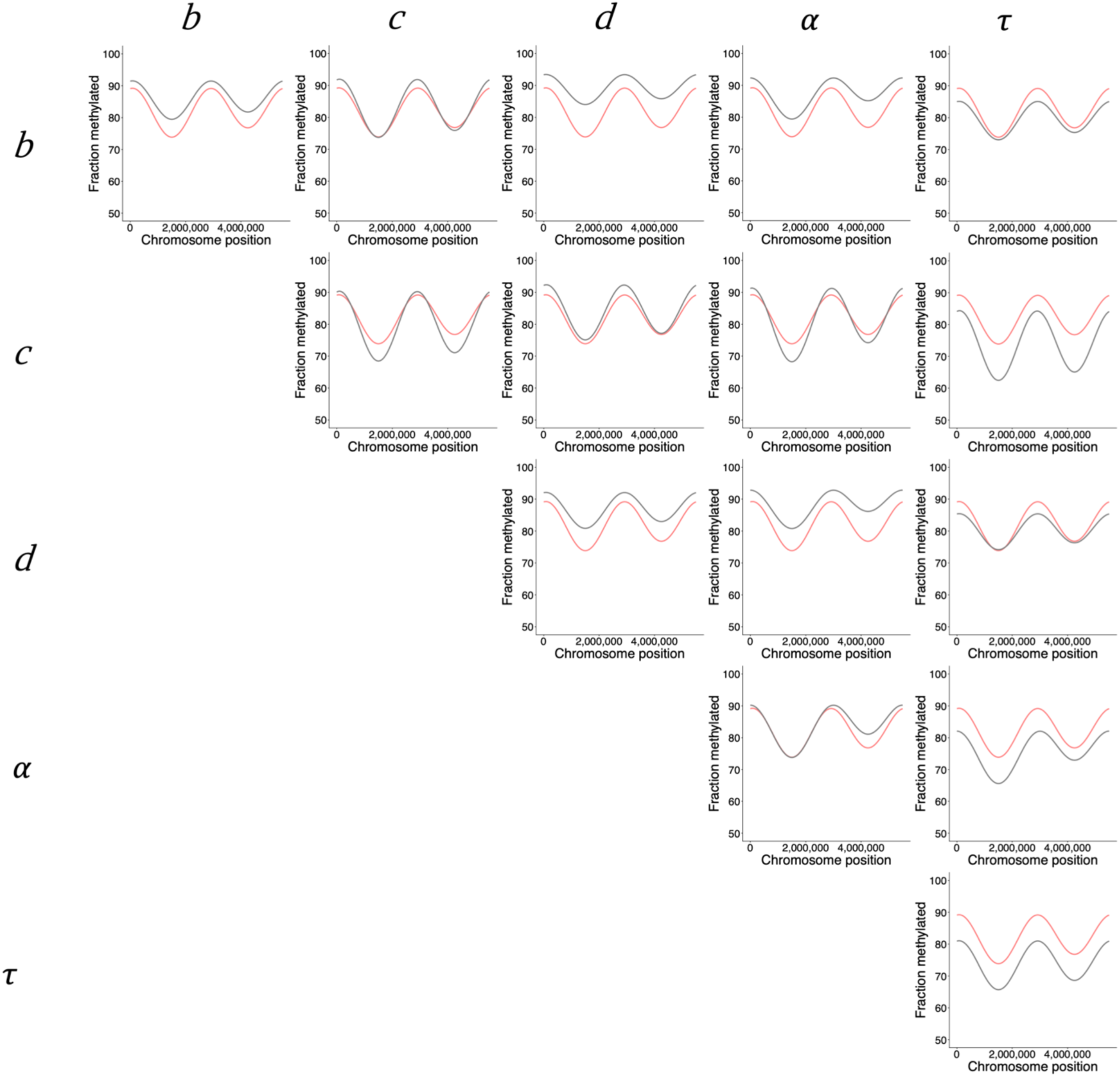
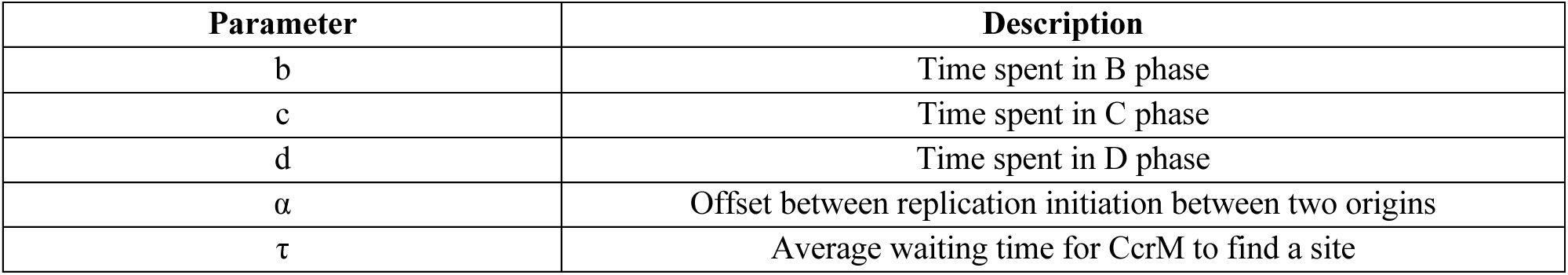
Effect of model parameters in shaping methylation patterns. In each panel, the gray curve represents the effect of changing the corresponding two parameters (on the row and the column) to 2x of their initial value used for the ‘null’ model – the red curve in all panels (same as Fig 1F). The diagonal entries simply represent the effect of changing just one parameter to 2x its initial value. Different parameters have distinct effects on the methylation patterns. Time spent in DNA replication (*c*) solely impacts the depth of the troughs, waiting time (*τ*) for CcrM activity decreases global methylation levels, offset between replication initiation between the two origins (*α*) impacts the asymmetry in methylation around the two peaks, and time spent in B and D phases (*b* and *d*) increases global methylation levels, primarily around the troughs.

**Fig S13.**
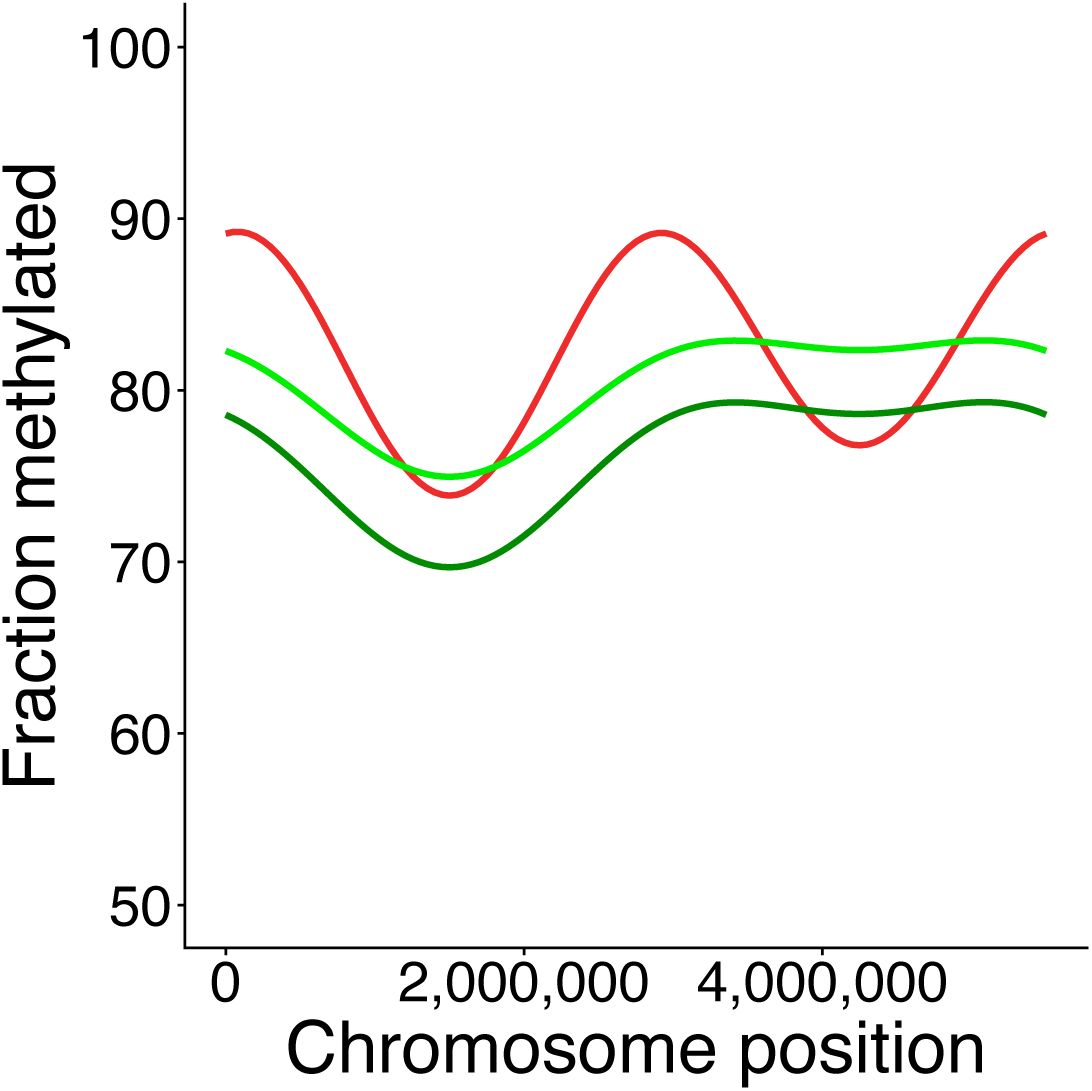
Methylome pattern predicted for increasing concentrations of ciprofloxacin stress. (green). Red curve represents the no stress prediction (same as panel Fig 3). For ciprofloxacin, we postulate an increase in time in D phase where the cell prepares to divide, presumably to mitigate DNA damage accrued during replication, and an increase in *α* – the offset in replication initiation between the two origins.

**Table S1.** Methylated motifs identified in *Methylobacterium extorquens* PA1 via nanopore sequencing. List of identified methylation motifs, the type of methylation, number of motifs in the genome (across both strands), and fraction of motifs in intergenic vs coding regions (16.07% of the genome is noncoding DNA). Underlined bases in the Motif column are the methylated bases. N denotes any of the 4 nucleotides.

| Motif | Methylation | Number | % Intergenic | % Coding | Gene |
| --- | --- | --- | --- | --- | --- |
| G<u>A</u>NTC | 6mA | 17090 | 23.8 | 76.2 | Mext_3181 |
| GCCG<u>A</u>TC | 6mA | 5591 | 10.2 | 89.8 | Mext_0272 |

**Table S2.** Strains used for comparing bias in distribution of methylation motifs in Fig S1.

| Strain | Group | NCBI RefSeq assembly |
| --- | --- | --- |
| *Methylobacterium extorquens* PA1 | Alphaproteobacteria | GCF_000018845.1 |
| *Methylobacterium nodulans* ORS2060 | Alphaproteobacteria | GCF_000022085.1 |
| *Methylobacterium phyllosphaerae* CBMB27 | Alphaproteobacteria | GCF_001936175.1 |
| *Methylobacterium radiotolerans* JCM 2831 | Alphaproteobacteria | GCF_000019725.1 |
| *Methylobacterium populi* BJ001 | Alphaproteobacteria | GCF_000019945.1 |
| *Methylobacterium extorquens* AM1 | Alphaproteobacteria | GCF_000022685.1 |
| *Sinorhizobium meliloti* MABNR56 | Alphaproteobacteria | GCF_037023865.1 |
| *Rhodopseudomonas palustris* RCB100 | Alphaproteobacteria | GCF_016584445.1 |
| *Mesorhizobium huakuii* NZP2235 | Alphaproteobacteria | GCF_034421895.1 |
| *Cereibacter sphaeroides* 2.4.1 | Alphaproteobacteria | GCF_003324715.1 |
| *Caulobacter crescentus* NA1000 | Alphaproteobacteria | GCF_000022005.1 |
| *Brucella abortus* 544 | Alphaproteobacteria | GCF_000369945.1 |
| *Agrobacterium tumefaciens* C58 | Alphaproteobacteria | GCF_000092025.1 |
| *Brevundimonas subvibrioides* ATCC 15264 | Alphaproteobacteria | GCF_000144605.1 |
| *Escherichia coli* MG1655 | Gammaproteobacteria | GCF_000005845.2 |
| *Salmonella enterica* LT2 | Gammaproteobacteria | GCF_000006945.2 |
| *Vibrio cholerae* E7946 | Gammaproteobacteria | GCF_013085165.1 |

**Table S3.** Hypomethylated GANTC sites during growth on succinate. . Using nanopore sequencing, we identify sites where the fraction of methylated reads for that site was less than 30% during exponential growth on succinate.

| Position | Context | Fraction methylated (%) |
| --- | --- | --- |
| 7198 | Intergenic (Mext 0007 ← / → Mext 0008) | 0.88 |
| 40637 | Within Mext 0036 | 22.7 |
| 144013 | Intergenic (→ Mext 0133) | 22.1 |
| 157138 | Intergenic (Mext 0141 ← / → Mext 0142) | 23.2 |
| 519613 | Within Mext 0474 | 23.4 |
| 519846 | Within Mext 0474 | 27.0 |
| 577609 | Intergenic (Mext 0529 ← / → Mext 0530) | 16.4 |
| 583166 | Intergenic (Mext 0534 ←) | 9.7 |
| 811556 | Within Mext 0746 | 6.1 |
| 811940 | Intergenic (→ Mext 0747) | 23.1 |
| 813036 | Intergenic (→ Mext 0747) | 15.6 |
| 1016685 | Intergenic (Mext 0939 → / ← Mext 0940) | 6.7 |
| 1048766 | Intergenic (Mext 0968 ← / → Mext 0969) | 26.3 |
| 1612376 | Within Mext 1439 | 29.6 |
| 2070524 | Within Mext 1852 | 29.9 |
| 2088251 | Within Mext 1865 | 6.6 |
| 2088683 | Within Mext 1865 | 27.7 |
| 2148927 | Intergenic (Mext 1931 ← / → Mext 1932) | 13.2 |
| 2738560 | Intergenic (Mext 2434 ← / → Mext 2435) | 29.9 |
| 2941475 | Intergenic (→ Mext 2626) | 21.8 |
| 3962426 | Intergenic (Mext 3580 ←) | 13.6 |
| 3989623 | Intergenic (→ Mext 3608) | 19.2 |
| 3990046 | Within Mext 3608 | 19.9 |
| 4050885 | Intergenic (→ Mext 3663) | 1.6 |
| 4050901 | Intergenic (→ Mext 3663) | 10.8 |
| 4326298 | Within Mext 3893 | 24.5 |
| 4400363 | Intergenic (Mext 3959 ← / → Mext 3960) | 22.1 |
| 4456706 | Intergenic (Mext 4012 ← / → Mext 4013) | 0 |
| 4833583 | Intergenic (Mext 4434 ← / → Mext 4435) | 25.6 |
| 4833591 | Intergenic (Mext 4434 ← / → Mext 4435) | 3.8 |
| 5199617 | Intergenic (Mext 4647 ← / → Mext 4648) | 14.2 |
| 5455782 | Intergenic (Mext 4880 → / ← Mext 4881) | 5.9 |
| 5459334 | Within Mext 4884 | 21.1 |

**Table S4.**
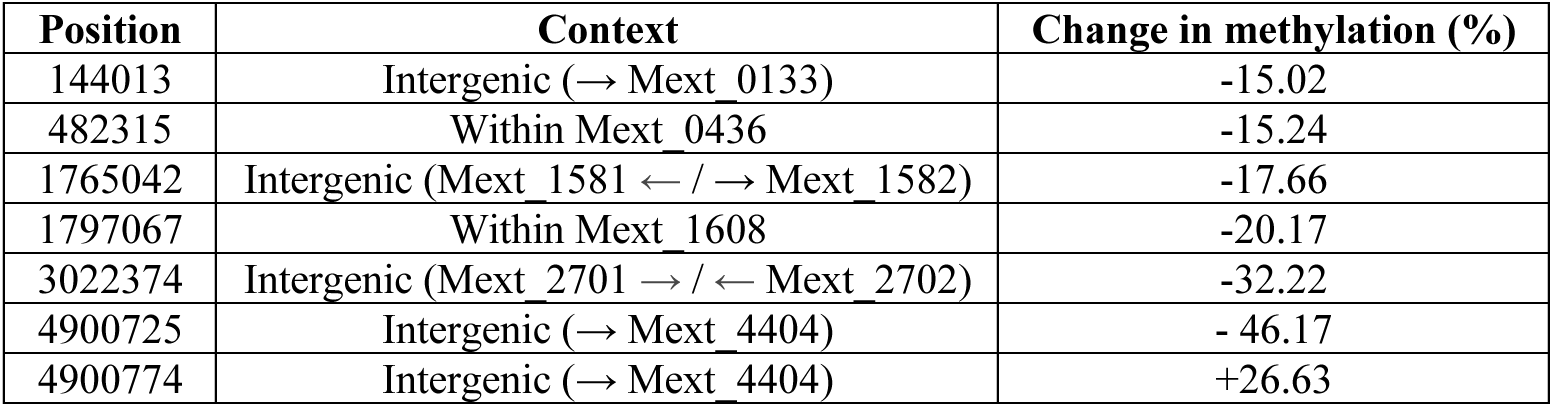
GANTC sites which show statistically significant changes in fraction of methylated reads during methanol growth compared to succinate. The location of the site on the chromosome, genetic context, and strength of change in methylation pattern is noted for all significant sites. Change in methylation pattern is classified as significant when *p* < 0.05 (Methods).

**Table S5.**
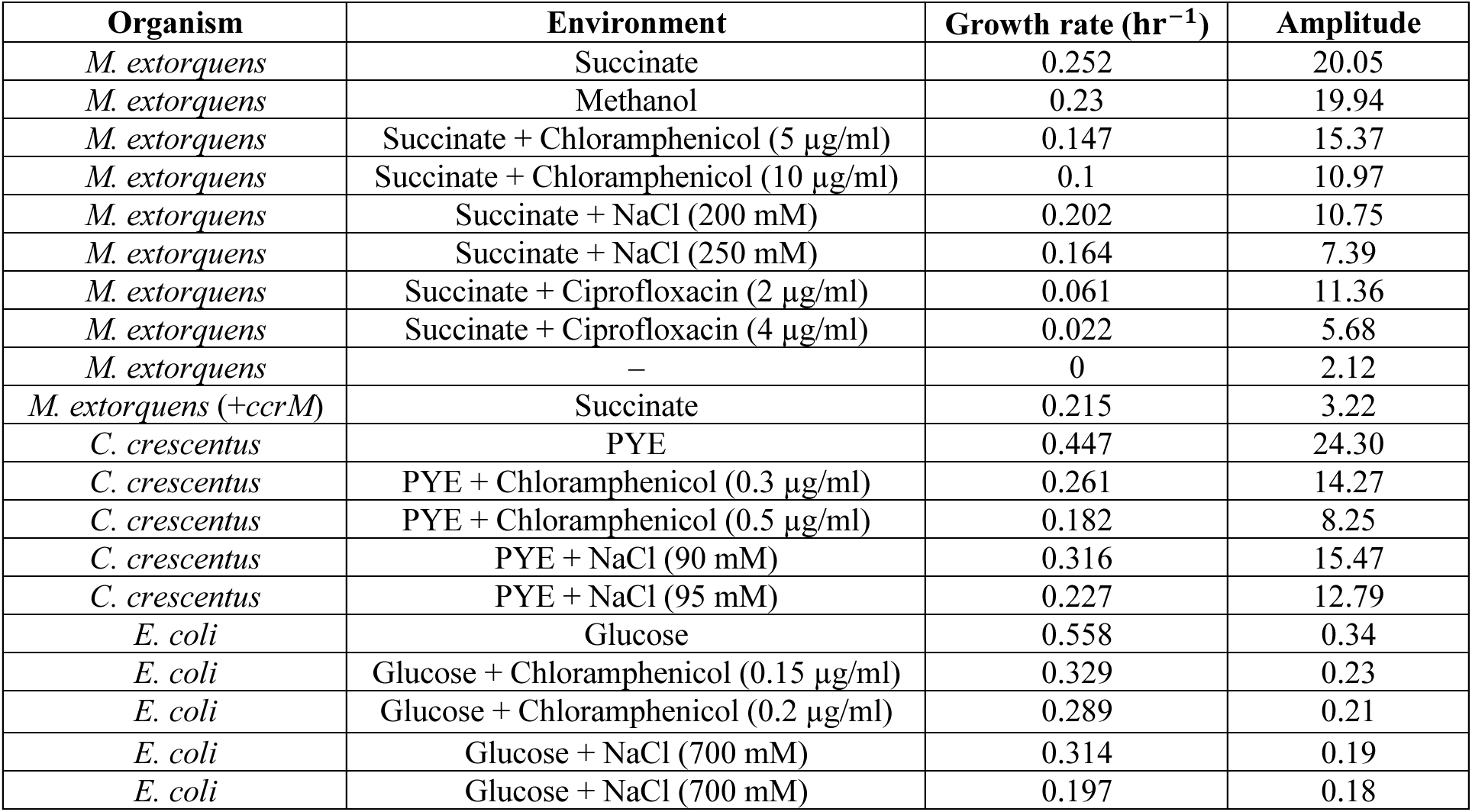
List of environments used to quantify growth rate and strength of chromosome position dependent methylation pattern in. **Fig 3F**. For *M. extorquens*, Mpipes minimal media was used throughout (Methods). For *E. coli*, M9 minimal media was used throughout (Methods). *For C. crescentus*, PYE was used as the base, and no additional carbon source was added.

**Table S6.** Model parameters used. (to characterize the methylome of unstressed cells in Fig S12). For testing the role of each parameter in shaping the methylation pattern, each variable was perturbed to 2x the value listed above and the effect on methylation noted (Fig S13).

| Parameter | Biological interpretation | Value (hrs) |
| --- | --- | --- |
| $b$ | Time spent in B phase | 1 |
| $c$ | Time needed for C phase (DNA replication) | 1 |
| $d$ | Time spent in D phase | 0.75 |
| $\alpha$ | Offset between replication initiation for the two origins | 0.1 |
| $\tau$ | Average waiting time for CcrM to find an unmethylated site | 0.5 |

**Table S7.**
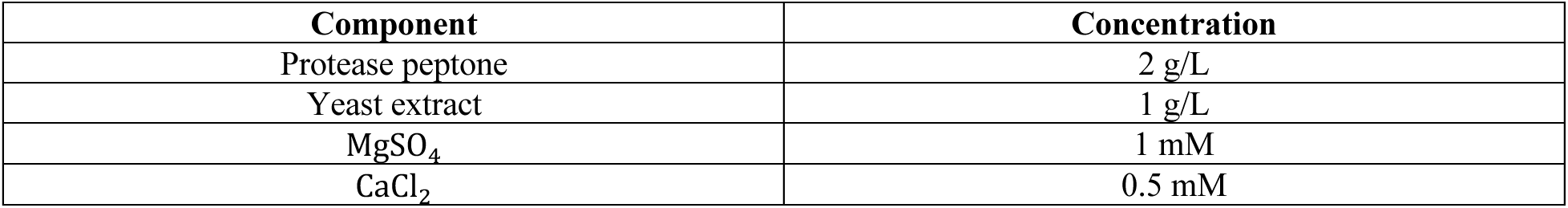
Components of the peptone yeast extract media used for *Caulobacter crescentus* growth.

| Component | Concentration |
| --- | --- |
| Protease peptone | 2 g/L |
| Yeast extract | 1 g/L |
| MgSO<sub>4</sub> | 1 mM |
| CaCl<sub>2</sub> | 0.5 mM |

## Notes

### Competing Interest Statement

The authors have declared no competing interest.

